# Neural mechanisms of postural sway-related beta-band oscillations: a cortico-basal ganglia-thalamic network model of intermittent control

**DOI:** 10.64898/2026.04.14.718368

**Authors:** Shota Tsugaya, Akihiro Nakamura, Taishin Nomura

## Abstract

Electroencephalographic (EEG) studies of human quiet stance demonstrate beta-band event-related desynchronization (beta-ERD) during the micro-fall phase of postural sway, followed by event-related synchronization (beta-ERS; post-movement beta rebound) during the micro-recovery phase. These modulations correlate with intermittent ankle muscle inactivation that exploits the stable manifolds of an unstable upright equilibrium; however, the underlying neurocircuit mechanisms remain elusive. Here, we investigated this rhythmogenesis using an embodied spiking neural network model of the cortico-basal ganglia-thalamic (CBGT) circuitry integrated with a physical inverted pendulum. In this closed-loop system, continuous sensory feedback is integrated into the striatum, while the motor cortex executes decisions via drift-diffusion-like population competition, where the decision time (DT) represents the intermittent control-off period. We demonstrate that simulated cortical LFPs exhibit characteristic sway-phase-locked beta-ERD and beta-ERS exclusively when corticostriatal synaptic weights are functionally balanced to implement intermittent control. Conversely, forced-choice continuous control that ceaselessly generates feedback torque fails to replicate these modulations, sustaining flat network states devoid of control-off periods (DT). Structural dissections reveal that while sensory drive remains continuous, sway-phase-locked beta modulations are an emergent property generated fundamentally by bidirectional thalamocortical loops and the intrinsic dynamics of the GPe-STN pacemaker circuit. Our findings suggest that CBGT-mediated, sway-phase-locked beta activity serves as a hallmark of healthy intermittent motor selection. This computational framework provides a crucial bridge linking pathological alterations in basal ganglia dynamics and the loss of behavioral intermittency to the postural impairments observed in clinical populations such as Parkinson’s disease.

**Significance Statement:** During quiet standing, human cortical beta oscillations dynamically modulate in phase with postural sway. Although this phenomenon is heavily linked to healthy motor control, the underlying neural mechanisms have remained unknown. This study resolves this ambiguity by developing a closed-loop computational model combining physical body dynamics with a cortico-basal ganglia-thalamic network. We demonstrate that human-like cortical beta modulations naturally emerge as an architectural property under an intermittent control strategy, driven fundamentally by the GPe-STN pacemaker circuit. Crucially, transitioning to a continuous control regime completely abolishes these oscillations. This framework establishes a direct neurocomputational link between the loss of behavioral intermittency and the pathological degradation of beta-band dynamics observed in patients with Parkinson’s disease.

## Introduction

Electroencephalography (EEG) provides a crucial window into the neural computations underlying motor control. Frequency-specific modulations of ongoing cortical activity, particularly within the beta-band (13–30 Hz), are closely linked to motor preparation and execution (Pfurtscheller and Lopes da Silva 1999). Specifically, event-related desynchronization (ERD) occurs during action preparation and execution, followed by event-related synchronization (ERS), or “beta rebound,” during movement cessation or status quo maintenance (Pfurtscheller and Lopes da Silva 1999; Stančák et al. 2000; Neuper and Pfurtscheller 2001; Steriade 2006). These dynamics carry profound clinical significance; patients with Parkinson’s disease exhibit pathological beta oscillations and altered modulation patterns that directly correlate with severe motor deficits (Brown et al. 2001; Jenkinson and Brown 2011; Moisello et al. 2015; Wu et al. 2019).

The cortico-basal ganglia-thalamic (CBGT) circuit is considered the primary neuroanatomical substrate for generating these movement-related beta rhythms (Roopun et al. 2008, 2010; Singh 2018; Vinding et al. 2019). Animal models suggest that glutamatergic inputs to the subthalamic nucleus (STN) and its reciprocal interactions with the globus pallidus (GP) play a critical role in rhythmic amplification (Tachibana et al. 2011). Despite extensive research, the exact mechanisms governing beta modulation remain controversial, partly due to the limited resolution of scalp EEG (Cohen 2017; Ouyang et al. 2022). While computational models have variously attributed beta oscillations to the STN-GPe circuit or the striatum (Terman et al. 2002; Bevan et al. 2002; Holgado et al. 2010; McCarthy et al. 2011; Pavlides et al. 2015; Ouyang et al. 2022), recent work shows that strong pallidostriatal loops drive low-beta (10–15 Hz) oscillations, whereas strong recurrent STN-GPe or intra-GPe connections drive gamma (>40 Hz) oscillations (Lindi et al. 2024). Crucially, most conventional models focus solely on isolated neural dynamics, neglecting bidirectional, closed-loop interactions with the musculoskeletal system.

Human quiet stance offers a unique and powerful paradigm to study this bidirectional interaction. Recent empirical findings show that postural sway is accompanied by transient cortical beta-ERD and beta-ERS, corresponding to the micro-fall and micro-recovery phases, respectively (Nakamura et al. 2023). These cortical modulations coincide with discrete bursts of calf muscle activity (Loram et al. 2005a), suggesting that cortical interventions in automatic postural control operate via discrete, intermittent commands (Asai et al. 2009; Gawthrop et al. 2011; Nomura et al. 2022) rather than continuous modulations (Winter et al. 1998). This supports an “intermittent control” strategy that dynamically exploits the stable manifold of an unstable upright equilibrium (Bottaro et al. 2008; Asai et al. 2009). The involvement of the basal ganglia in this process is corroborated by recent work demonstrating that an intermittent control strategy—defined by switching boundaries in the state space—can be effectively acquired through reinforcement learning (Takazawa et al. 2024). Because the CBGT circuit is the primary neural substrate for reinforcement learning, postural stability may be maintained by optimizing synaptic efficacies within this loop to facilitate healthy, intermittent motor selection.

In this study, we investigated the neural mechanisms underlying sway-related beta oscillations using a spiking neural network (SNN) model of the CBGT circuitry. Crucially, the model stabilizes a physical inverted pendulum in real time, while simultaneously simulating EEG via the summation of cortical postsynaptic currents. By systematically adjusting the synaptic weights of cortico-striatal projections— key targets for motor learning and dopamine depletion (Cataldi et al. 2022)—we demonstrate that simulated EEG exhibits realistic beta-ERD and beta-ERS only when the network is functionally tuned for intermittent control operated under small feedback gains. Conversely, forceful control operated continuously under large feedback gains fails to produce these characteristic cortical modulations. While Lindi et al. (Lindi et al. 2024) demonstrated the generation mechanisms of parkinsonian beta oscillations within a disembodied network, our closed-loop neuro-biomechanical computational model uncovers the mechanistic origins of beta modulation during both healthy and pathological balance control. Ultimately, our results provide a novel computational link between the functional tuning of the basal ganglia and the emergence of cortical rhythms essential for behavioral postural stability.

## Materials and Methods

### Physiological Foundations and Abstractions of CBGT Circuit Topology

We developed a spiking neural network (SNN) model of the cortico-basal ganglia-thalamic (CBGT) circuit by adapting a previous neural mass model designed for simulating finger tapping (Ursino et al. 2020). In our framework, we replaced each neural mass component with a population of spiking neurons and integrated a physical model of an inverted pendulum, which was absent in the original Ursino model. Within this closed-loop architecture, sensory information regarding the pendulum’s kinematic state is represented by neuronal firing in the sensory cortex and fed back to the striatum. The motor cortex, positioned at the output terminal of the CBGT loop, decides between dorsiflexion and plantarflexion via competition between antagonistic motoneuronal populations. This competitive process follows drift-diffusion dynamics (Ratcliff et al. 2016), where the decision time (DT) represents the “control-off period” of the intermittent controller. Preliminary parameter exploration indicated that the synaptic weights (*W*) projecting from the sensory cortex to the striatum are the critical determinants governing the degree of competition between the two cortical motoneuron populations; we therefore tuned these parameters as detailed below.

We implemented variants of the CBGT model, including those corresponding to an intermittent control model operated under small feedback gains and forceful control model operated continuously under large feedback gains, where typical values of small and large gains were employed, respectively, based on the previous study that assimilates postural sway data from healthy people and patients with Parkinson’s disease with severe postural impairment (Suzuki et al. 2020). Using these models, we conducted two distinct computational experiments. Experiment 1 involved a postural control task to stabilize the inverted pendulum and reconstruct EEG dynamics during quiet stance. This experiment was designed to compare movement-related beta modulations between the intermittent CBGT model and the continuous stiffness control model. Experiment 2 was conducted to dissect the network mechanisms underlying the generation of movement-related beta modulations. In this experiment, the transmission line conveying motor commands from the cortex to the pendulum was disconnected, and the pendulum was moved passively along a prescribed, periodic trajectory independent of the CBGT cortical output. This setup allowed us to examine how model-generated EEG dynamics are influenced by state-dependent sensory signals under various network conditions, without confounding the analysis with the requirements of postural stability.

### Inverted pendulum model

To simulate the human body during quiet stance, we employed an inverted pendulum model, which simplifies a previously validated postural model (Asai et al. 2009). The inverted pendulum rotates in the sagittal plane around a pinned joint corresponding to the ankle. The dynamics of the system are governed by the following equation of motion:

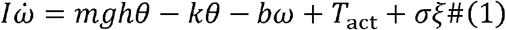

where *θ* and *ω* are the tilt angle and angular velocity of the pendulum, respectively; *I* (60 kgm^2^) is the moment of inertia of the pendulum around the joint; *m* (60 kg) is the pendulum mass; *g* (9.81 m/s^2^) is the gravitational acceleration; and *h*(1.0 m) is the distance from the joint to the center of mass. The parameters *k*(471 Nm/rad) and *b*(4.0 Nms/rad) represent the passive torsional elasticity and the viscosity of the joint, respectively, derived from empirical identifications (Loram and Lakie 2002; Casadio et al. 2005). The term *σ ξ* represents zero-mean Gaussian white noise with a standard deviation of *σ*. The term *T*_act_ represents the active feedback torque selected by the motor cortex of the CBGT model at each simulation time step as from three possible actions: {−|*Pθ*_Δ_ + *Dω*_Δ_|, 0,|*P θ*_Δ_+ *D ω*_Δ_|}, where *θ*_Δ=_ θ (*t* − Δ) and *ω*_Δ_=*ω* (*t* − Δ) represent the time-delayed state of the pendulum, reflecting a neural transmission delay of Δ = 0.2 s. The parameters *P* and *D* are the proportional and derivative feedback gains to generate backward torque (*T*_act_ = −|*P*_Δ_*θ*;+ *Dω*_Δ_ |)or forward torque (*T*_act_ =| *Pθ*_Δ_ + *Dω*_Δ_ |). A zero torque condition (*T*_act_=0) corresponds to the periods when the active feedback controller is switched off in the intermittent control model (Asai et al. 2009).

Action selection among these three choices is determined by the CBGT model as described below. For all simulations, the equation of motion and the equations governing spiking neural dynamics were discretized with a time step of *δt*=0.001 s and integrated numerically using the forward Euler-Maruyama method.

### CBGT circuit model

#### Network architecture

The CBGT circuit model was structured as illustrated in Fig. 1, following the topological framework of Ursino et al. (Ursino et al. 2020). The network processes the delayed physical state of the inverted pendulum and selects one of the three active torques (*T*_act_) to maintain upright stability. Concurrently, the spiking dynamics of cortical populations generate local field potentials (LFPs) to simulate cortical EEG.

**Figure 1:**
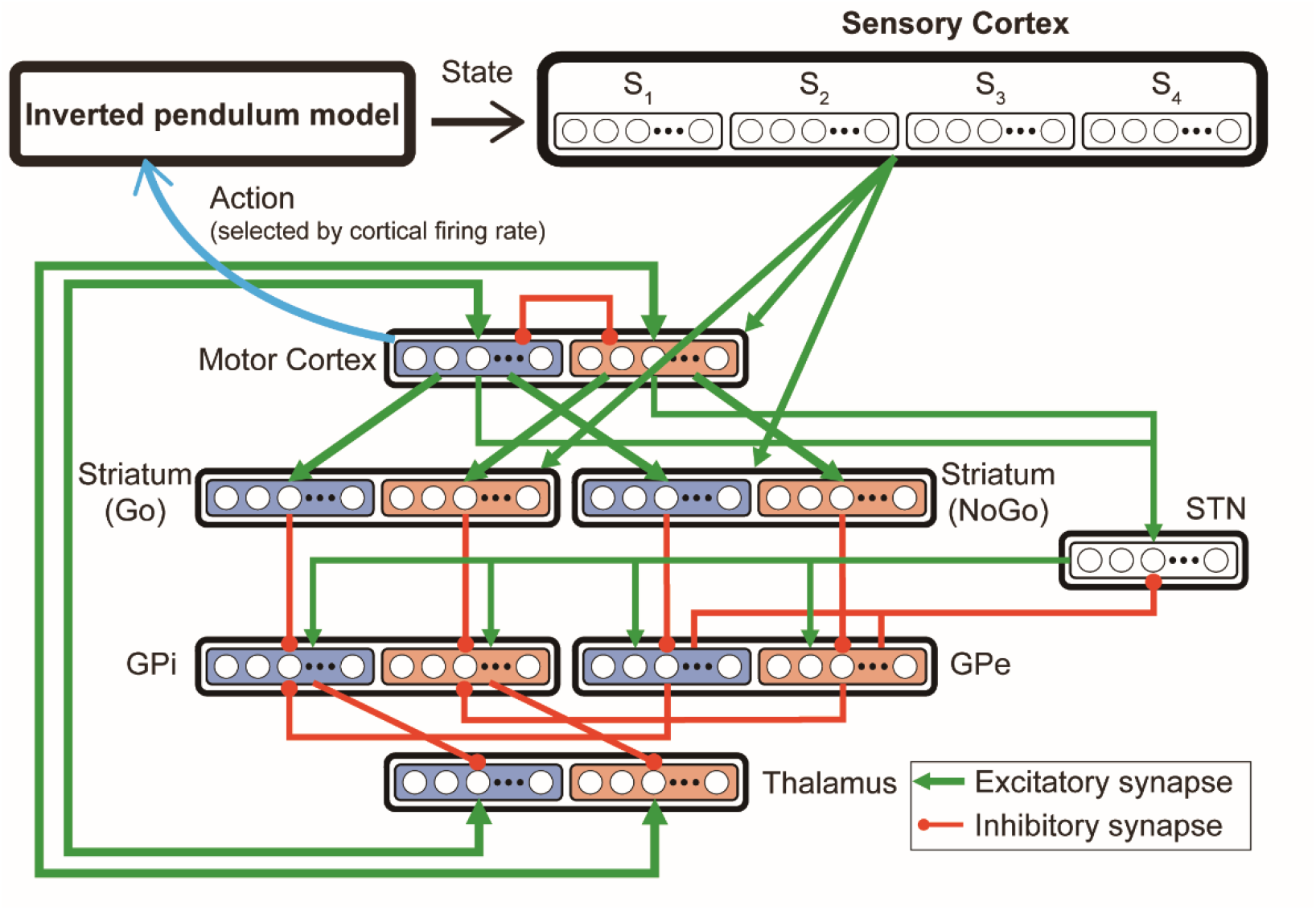
Schematic diagram of the network model for the cortico-basal ganglia thalamic (CBGT) loop developed in this study. This model consists of neuronal populations of sensory cortex, cortex, striatum, globus pallidus pars externa (GPe), globus pallidus pars interna (GPi), subthalamic nucleus (STN), and thalamus. Each neural population except STN population includes two populations (colored blue and orange). Blue and orange populations are associated with backward- and forward-torque actions, respectively. Green lines with arrowheads represent excitatory injection and red lines with circular-heads inhibitory projections, respectively. Specifically, reciprocal excitatory connections exist between the motor cortex and thalamus, where thalamic neurons project back to their corresponding agonistic motor cortical sub-populations (e.g., backward thalamus to backward motor cortex) to complete the thalamocortical feedback loop. In this model, the state of the inverted pendulum is transmitted to and represented in the sensory cortex, and active torque (either backward torque, forward torque, or null torque) is selected by the spike rate of two populations of the cortical neurons.

The model encompasses several interconnected neuronal populations: the sensory cortex, motor cortex, striatum, globus pallidus pars externa (GPe), globus pallidus pars interna (GPi), subthalamic nucleus (STN), and thalamus. The striatum is segregated into D1 receptor-expressing (Go-striatum, direct pathway) and D2 receptor-expressing (NoGo-striatum, indirect pathway) populations. Except for the STN, which comprises a single population of 500 neurons, each nucleus consists of 1,000 neurons split into two functionally distinct sub-populations of 500 neurons each: a “backward” population (colored blue in Fig. 1) and a “forward” population (colored orange in Fig. 1), corresponding to the actions for backward and forward torques, respectively. Neurons within the same sub-population lack intra-group connectivity but project to their target agonistic sub-populations in an all-to-all manner. For instance, cortical neurons in the backward population project exclusively to backward-related sub-populations in both the Go- and NoGo-striatum. Excitatory and inhibitory pathways are modeled via glutamatergic and GABAergic projections, respectively (Fig. 1). Specifically, reciprocal excitatory connections exist between the motor cortex and thalamus, where thalamic neurons project back to their corresponding agonistic motor cortical sub-populations (e.g., backward thalamus to backward motor cortex) to complete the thalamocortical feedback loop. The sensory cortex contains 2,000 neurons, whose firing rates encode the delayed state variables (*θ*_Δ_ and *ω*_Δ_) and drive downstream cortical and striatal populations.

In this modeling of embodied posture control, the spiking CBGT network incorporated both established neuroanatomy and functionally principled computational abstractions. In accordance with the known parallel-channel organization of the cortico-basal ganglia loop (Alexander and DeLong 1985; Hoover and Strick 1993), neurons in each CBGT subregion (motor and sensory cortices, striatum, GPe, STN, GPi, thalamus) are segregated into two distinct, non-overlapping populations governing opposing movement directions: forward-torque channel and backward-torque channel. This dual-channel design directly mirrors the antagonist muscle pair acting at the ankle joint during quiet standing—namely, plantarflexors (Soleus and Gastrocnemius) resisting forward sway, and dorsiflexors (Tibialis Anterior) resisting backward sway. The anatomical separation of antagonist muscle representations in primary motor cortex (Rathelot and Strick 2006) and the ability of striatal microstimulation to elicit single-joint directional movements (Alexander and DeLong 1985; Nambu et al. 2002) provide strong neurophysiological justification for this channel segregation. For sensory state encoding, a mapping of a biomechanical state onto firing of spatially distributed neurons of sensory cortex serves as a functional abstraction representing multi-sensory integration (somatosensory, vestibular, and visual) conveyed to the striatum via sensory cortical inputs. Rather than modeling the full sensory cortical population, state-dependent corticostriatal projection weights *W* represent the functional tuning of sensory-to-striatal synaptic strength, encoding contextual state-action mappings across Go (Direct) and NoGo (Indirect) striatal spiny projection neurons.

#### Neuron model

Individual neurons are modeled as leaky integrate-and-fire (LIF) units without detailed biophysical ion-channel dynamics. The membrane potential integrates synaptic currents over time and fires an action potential upon reaching a fixed threshold. The LIF model accounts for a leak current that drives the potential toward its resting state (Izhikevich 2004). The membrane potential *V* of a single neuron, as the time-dependent dynamic variable, evolves as follows (Ouyang et al. 2022):

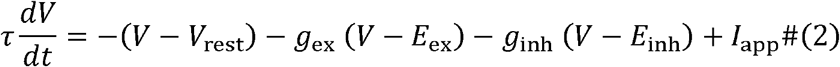

where *τ* = 20 ms is the membrane time constant, *V*_rest_ = −60 mV is the resting potential, *E*_ex_ = 0 mV is the excitatory reversal potential, and *E*_inh_ = −80 mV is the inhibitory reversal potential. The term *I*_app_ represents an external input current composed of a population-specific baseline constant (Table 1) and additive, independent Gaussian white noise. These population-specific constants reflect physiological background drives; for example, striatum neurons receive the dopaminergic bias (*D*=1.0), while GPe and GPi neurons receive constant current to sustain spontaneous baseline firing.

**Table 1.** Parameters of the external input current *I*_app_, in a LIF neuron model. Striatum (Go and NoGo) neurons receive dopaminergic input (*I*_app_ =*α* · *D* with *α* = 3.25 for Go neurons and *I*_app_ =*β* · D with *β* = −3.9 for NoGo neurons, D=1.0 in this study). Unlike the Ursino model, external inputs to Go neurons were assumed to be always excitatory input for simplicity. NoGo neurons receive inhibitory input. Thus, the constant term (external input current *I*_app_) is always positive in Go neurons and negative for NoGo neurons. GPe (globus pallidus pars externa) and GPi (globus pallidus pars interna) receive excitatory background input to exhibit certain basal activity.

| Neuron population | Value of the constant term |
| --- | --- |
| Go | 3.25 |
| NoGo | -3.9 |
| GPe | 18 |
| GPi | 24 |

The time-dependent excitatory synaptic conductance (*g*_ex_) for a given postsynaptic neuron is defined as:

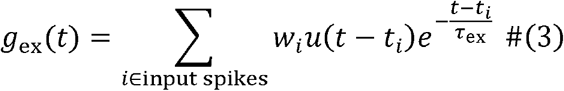

where *u*(*t*) is the Heaviside step function (*u*(*t*) =1 for *t* ≥0, and 0 otherwise). The coefficient *w*_*i*_ defines the synaptic coupling strength from the presynaptic neuron delivering the *i*-th spike at time *t*_*i*_. These coupling weights depend on the identity of the presynaptic and postsynaptic populations (see Tables 2, 3 and 4). To introduce structural heterogeneity, the 500×500 connectivity matrix between populations was generated by scaling random numbers drawn from a Cauchy distribution by a population-specific coefficient (Tables 2, 3 and 4). The excitatory decay constant was set to *τ*_ex_ = 5 ms. The time-dependent inhibitory synaptic conductance (*g*_inh_) was modeled analogously, with a decay constant of τ_inh_ = 10 ms.

**Table 2.** Parameters of the synaptic weight in the present network model, which are common for the intermittent and continuous CBGT controller. The synaptic weight index of size 500×500 is generated by multiplying a random number that follows a Cauchy distribution by the coefficient values in this table.

| Presynaptic neuron | Postsynaptic neuron | Synapse type | Coefficient name and value |
| --- | --- | --- | --- |
| Sensory Cortex | Cortex | Excitatory | $W_{S2C} \quad 7.5 \times 10^{-3}$ |
| Cortex | Cortex | Inhibitory | $W_{LAT} \quad 7.5 \times 10^{-2}$ |
| Cortex | Go | Excitatory | $W_{C2G} \quad 2.54 \times 10^{-3}$ |
| Cortex | NoGo | Excitatory | $W_{C2N} \quad 4.475 \times 10^{-3}$ |
| Cortex | STN | Excitatory | $W_{C2STN} \quad 3.00 \times 10^{-2}$ |
| Cortex | Thalamus | Excitatory | $W_{C2T} \quad 4.655 \times 10^{-2}$ |
| Go | GPI | Inhibitory | $W_{G2I} \quad 4.35 \times 10^{-2}$ |
| NoGo | GPe | Inhibitory | $W_{N2E} \quad 2.6 \times 10^{-2}$ |
| GPe | GPI | Inhibitory | $W_{E2I} \quad 3.21 \times 10^{-2}$ |
| GPe | STN | Inhibitory | $W_{E2STN} \quad 2.085 \times 10^{-2}$ |
| GPI | Thalamus | Inhibitory | $W_{I2T} \quad 2.875 \times 10^{-2}$ |
| STN | GPe | Excitatory | $W_{STN2E} \quad 2.85 \times 10^{-2}$ |
| STN | GPI | Excitatory | $W_{STN2I} \quad 5.345 \times 10^{-2}$ |
| Thalamus | Cortex | Excitatory | $W_{T2C} \quad 9.64 \times 10^{-3}$ |

**Table 3.** Parameters of the synaptic weight from the sensory cortex to the striatum in the intermittent CBGT model.

| State | Neuronal population | Action-related Population | Coefficient name and value |  |
| --- | --- | --- | --- | --- |
| S<sub>1</sub> | Go | Forward torque | G<sub>min</sub> | 3.0×10<sup>-3</sup> |
|  |  | Backward torque | G<sub>max</sub> | 1.0×10<sup>-2</sup> |
|  | NoGo | Forward torque | N<sub>max</sub> | 1.7×10<sup>-2</sup> |
|  |  | Backward torque | N<sub>min</sub> | 5.0×10<sup>-3</sup> |
| S<sub>2</sub> | Go | Forward torque | W<sub>S2G</sub> | 6.5×10<sup>-3</sup> |
|  |  | Backward torque | W<sub>S2G</sub> | 6.5×10<sup>-3</sup> |
|  | NoGo | Forward torque | W<sub>S2N</sub> | 1.1×10<sup>-2</sup> |
|  |  | Backward torque | W<sub>S2N</sub> | 1.1×10<sup>-2</sup> |
| S<sub>3</sub> | Go | Forward torque | G<sub>max</sub> | 1.0×10<sup>-2</sup> |
|  |  | Backward torque | G<sub>min</sub> | 3.0×10<sup>-3</sup> |
|  | NoGo | Forward torque | N<sub>min</sub> | 5.0×10<sup>-3</sup> |
|  |  | Backward torque | N<sub>max</sub> | 1.7×10<sup>-2</sup> |
| S<sub>4</sub> | Go | Forward torque | W<sub>S2G</sub> | 6.5×10<sup>-3</sup> |
|  |  | Backward torque | W<sub>S2G</sub> | 6.5×10<sup>-3</sup> |
|  | NoGo | Forward torque | W<sub>S2N</sub> | 1.1×10<sup>-2</sup> |
|  |  | Backward torque | W<sub>S2N</sub> | 1.1×10<sup>-2</sup> |

**Table 4.**
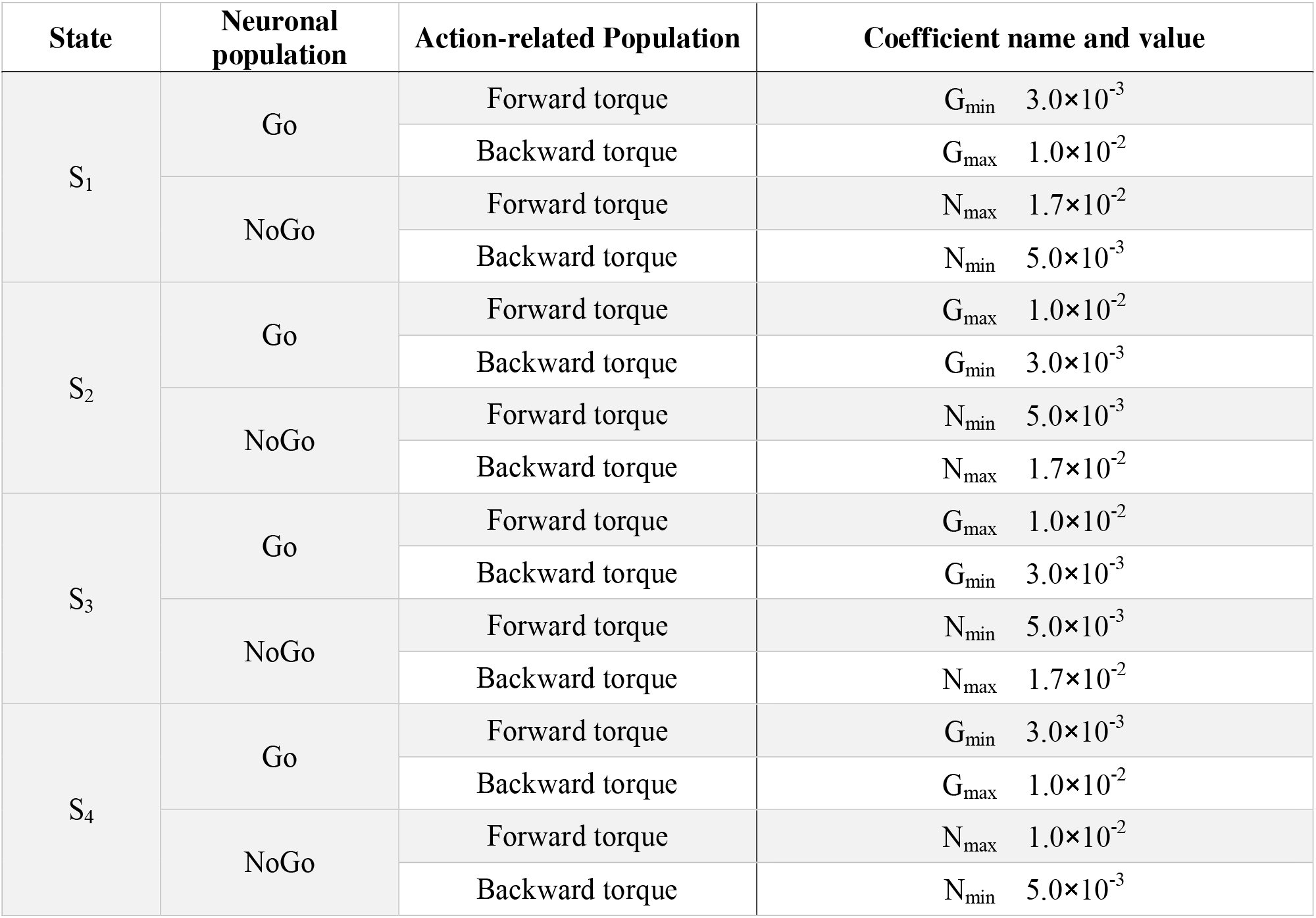
Parameters of the synaptic weight from the sensory cortex to the striatum in the forced-choice CBGT model.

| State | Neuronal population | Action-related Population | Coefficient name and value |  |
| --- | --- | --- | --- | --- |
| S<sub>1</sub> | Go | Forward torque | G<sub>min</sub> | 3.0×10<sup>-3</sup> |
|  |  | Backward torque | G<sub>max</sub> | 1.0×10<sup>-2</sup> |
|  | NoGo | Forward torque | N<sub>max</sub> | 1.7×10<sup>-2</sup> |
|  |  | Backward torque | N<sub>min</sub> | 5.0×10<sup>-3</sup> |
| S<sub>2</sub> | Go | Forward torque | G<sub>max</sub> | 1.0×10<sup>-2</sup> |
|  |  | Backward torque | G<sub>min</sub> | 3.0×10<sup>-3</sup> |
|  | NoGo | Forward torque | N<sub>min</sub> | 5.0×10<sup>-3</sup> |
|  |  | Backward torque | N<sub>max</sub> | 1.7×10<sup>-2</sup> |
| S<sub>3</sub> | Go | Forward torque | G<sub>max</sub> | 1.0×10<sup>-2</sup> |
|  |  | Backward torque | G<sub>min</sub> | 3.0×10<sup>-3</sup> |
|  | NoGo | Forward torque | N<sub>min</sub> | 5.0×10<sup>-3</sup> |
|  |  | Backward torque | N<sub>max</sub> | 1.7×10<sup>-2</sup> |
| S<sub>4</sub> | Go | Forward torque | G<sub>min</sub> | 3.0×10<sup>-3</sup> |
|  |  | Backward torque | G<sub>max</sub> | 1.0×10<sup>-2</sup> |
|  | NoGo | Forward torque | N<sub>max</sub> | 1.0×10<sup>-2</sup> |
|  |  | Backward torque | N<sub>min</sub> | 5.0×10<sup>-3</sup> |

#### Physical state representation in the sensory cortex

For computational simplicity, the delayed physical state of the pendulum (*θ*_Δ_, *ω*_Δ_) is categorized into four state-dependent regions (S_1_-S_4_) within the phase plane, corresponding roughly to the four quadrants (Fig. 2). Specifically, the phase space is partitioned by the lines ω_Δ_ = −(*P/D*) θ_Δ_ and the *θ*-axis into the following four regions:

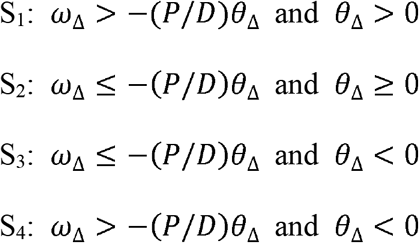

**Figure 2:**
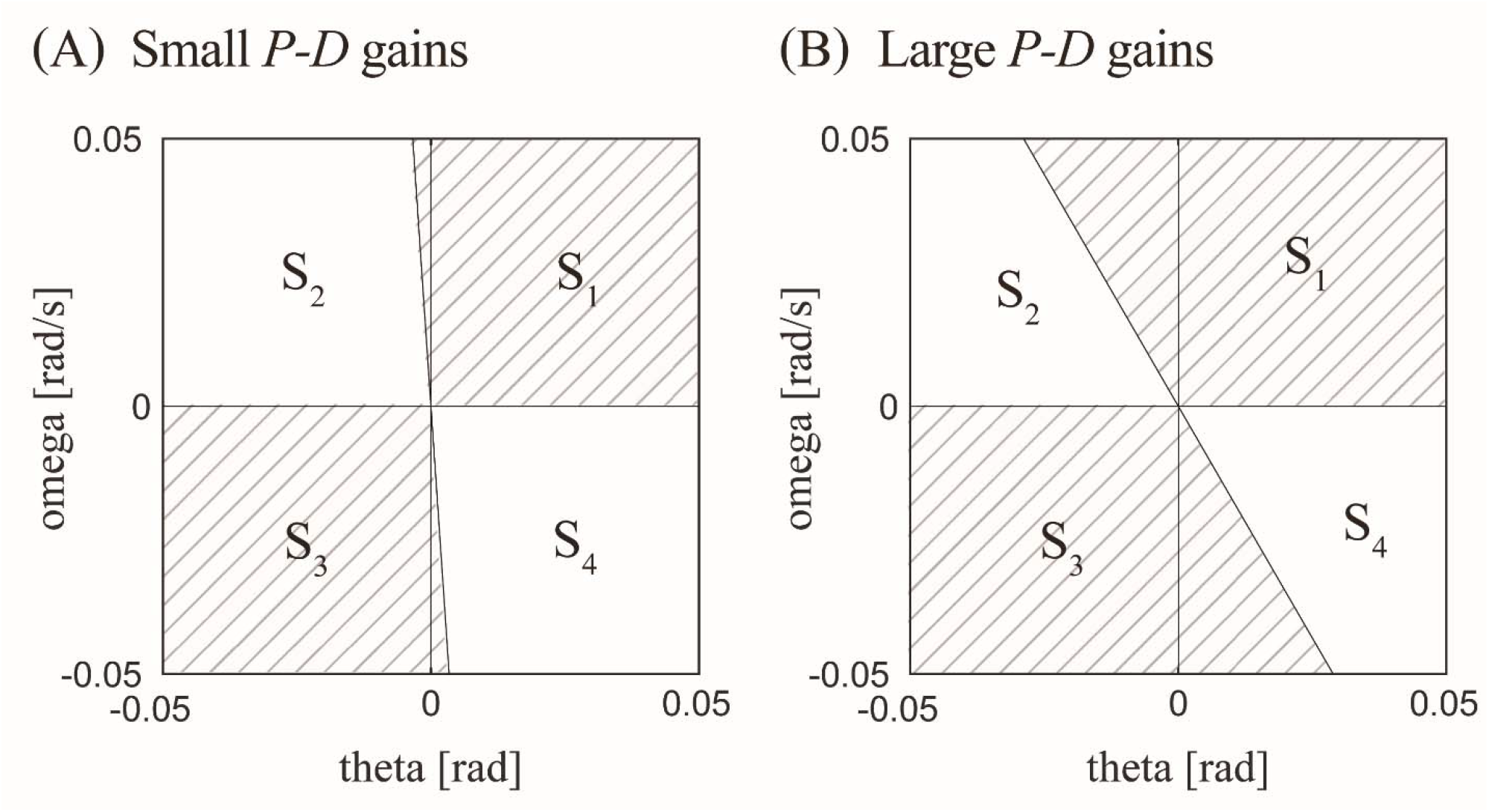
Categorical representation and encoding of the physical state of the inverted pendulum based on the position of the state point in the phase plane. The state of inverted pendulum is encoded based on the four regions (*S*_1_, *S*_2_, *S*_3_, *S*_4_) in the phase plane. The negatively tilted line through the origin is defined as *ω* = −(*P/D*) *θ*, where (*P,D*) = (0.*45mgh*, 18) in (A)with the small *P*-*D* gains and (*P,D*) = 0.5*mgh*, 150) in (B) with the large *P*-*D* gains.

The 2,000 sensory cortical neurons are divided into four identical pools of 500 neurons, each dedicated to one of these regions. When the delayed state resides within region S_*i*_ (*i*=1,2,3,4), the corresponding pool of 500 neurons fires stochastically according to an independent Poisson process, thereby mapping the system state to sensory firing rates. The probability of a single neuron generating a spike within a time step *δt=*0.001 s is given by (Johnson 1996):

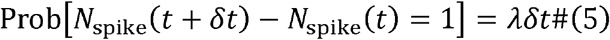

where *N*_spike_ (*t*) is the cumulative number of firings for a neuron until time *t*, and λ is the constant Poisson intensity (firing rate). We set λ = 100 Hz for neurons assigned to the active region S_*i*_, and λ= 0 Hz otherwise. In Experiment 2, we also tested λ = 200 Hz to assess how sensory signal intensity affects the simulated EEG.

#### Implementation of the intermittent and forced-choice continuous-like CBGT controllers

We configured the network parameters to realize two different control regimes for postural stabilization: an intermittent controller and a forced-choice continuous-like controller. These regimes were implemented by systematically altering the synaptic weights *W* for the sensorimotor cortico-striatal projections. As mentioned above, we employed two sets of typical *P*-*D* feedback gains and noise intensity (*σ*): a small gains-noise set and a large gain-noise set, respectively, based on the previous study that assimilates postural sway data from healthy people and patients with Parkinson’s disease with severe postural impairment (Suzuki et al. 2020).

#### Intermittent CBGT controller

In the intermittent control framework, the active feedback controller is switched off when the state point of the pendulum moves near the stable manifold of the unstable, saddle-type upright equilibrium (Asai et al. 2009). Regions S_2_ and S_4_ represent these “safe” zones, where the passive dynamics transiently pull the pendulum back toward the vertical position without active intervention. Conversely, the active feedback controller is switched on when the state point of the pendulum enters regions S_1_ or S_3_, which lie near the unstable manifold.

The motor cortex evaluates and selects the appropriate motor command through firing rate competition between spike rates of the backward and forward populations. Sensory signals generated in the sensory cortex drive this loop by stimulating the corresponding Go or NoGo striatal sub-populations. These signals propagate through the GPe, GPi, and STN to the thalamus, ultimately feeding back to the cortex via thalamocortical projections. Excitatory postsynaptic currents from these projections accumulate in the cortical neurons. Firing rate competition operates between the aggregate spike counts per second of the backward and forward cortical populations. To ensure high temporal resolution in action selection, firing rate and motor commands are evaluated and updated continuously at every simulation time step (δ*t* = 0.1ms). That is, at each time step *t*, the population firing rate of the motor cortex is computed using a 0.1-s moving time window ([*t* − 0.1 *s,t*.]) If this smoothed firing rate exceeds the decision threshold, an active feedback torque is exerted on the ankle joint. Thus, the temporal resolution of the action selection process matches the numerical integration step, ensuring that the observed intermittency in torque generation arises dynamically from the network’s spiking threshold dynamics rather than from discrete temporal sampling.

More specifically, in the context of a drift-diffusion model, an increase in a population’s firing rate represents the accumulation of evidence toward its corresponding action. If the firing rate of the backward population reaches the decision threshold (50 Hz) first, the backward torque (*T*_act_ = −| *Pθ*_Δ_+ *D ω*_Δ_ |) is applied to the pendulum. Conversely, if the forward population reaches the threshold first, the forward torque (*T*_act_ − | *Pθ*_Δ_+ *D ω*_Δ_|) is executed. In this study, the threshold for active torque selection was set at 50 Hz throughout the study. The threshold was fixed at 50 Hz based on established studies (Mohanty et al. 2025). The selected torque remains active as long as the winning population’s firing rate stays above the threshold. A period during which both populations remain below the 50 Hz threshold defines the decision time (DT), during which no active torque is applied (*T*_act_ − 0), implementing the “control-off” phase of the intermittent strategy. Reciprocal inhibition between the two cortical populations enforces a winner-take-all dynamic (Ursino et al. 2020) (Fig. 1).

Because sensory cortico-striatal weights *W* govern the drift bias in this competition, we explored these connections to satisfy the intermittent control criteria. More specifically, we set the small *P*-*D* gains (and noise intensity) in the model, by which upright posture of the inverted pendulum cannot be stabilized, if the small-gain delay PD feedback controller operates persistently, due to delay-induced instability. Thus, the exploration was performed to find *W* that could stabilize the upright posture, not to reproduce beta modulations in the simulated EEG, which was theoretically equivalent to finding *W* that could realize intermittent control. In this way, the appropriate weights *W* realize the following characteristics. When the pendulum enters S_1_ (tilted forward and falling), the backward and forward population of Go-striatum receives strong and weak inputs, respectively, ensuring a high probability of selecting backward torque. Concurrently, the backward and forward populations of the NoGo-striatum receive weak and strong inputs, respectively, preventing the antagonistic action from interfering with the command (Fig. 3A-1, Table 3A). Conversely, when the pendulum is in S_3_ (tilted backward and falling), the forward Go-striatum and the backward NoGo-striatum receives strong inputs, while their counterparts receive weak inputs, favoring forward torque (Fig. 3A-3, Table 3A). Crucially, when the trajectory enters the safe regions (S_2_ or S_4_), sensory inputs to the antagonistic striatal populations are balanced with equal strength. This unbiased projection ensures close, prolonged competition between the populations (Fig. 3A-2 and Fig. 3A-4; Table 3A), extending the DT and maximizing the probability of selecting the null action (*T*_act_ = 0). The sensoristriatal synaptic weight vector *W* that satisfied the characteristics detailed here is referred to as *W*_unbiased_, which leads to balanced, unbiased action selections.

**Figure 3:**
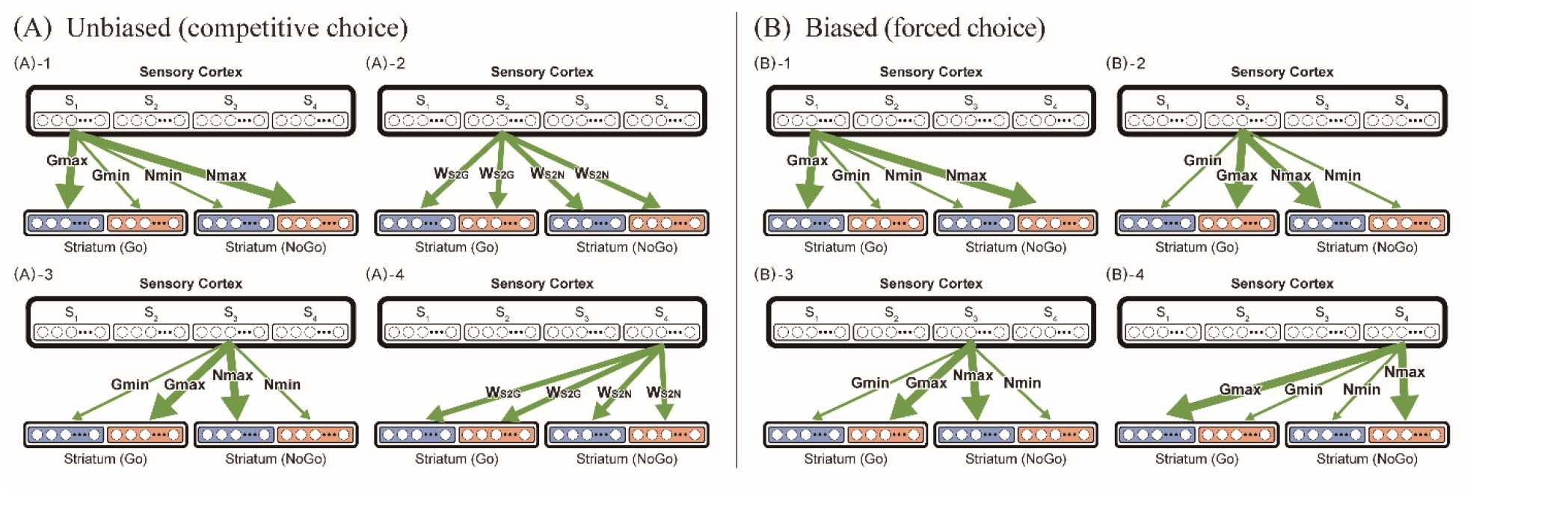
The state-dependent synaptic weights of the sensory cortico-striatal projections for the intermittent and forced-choice control model. (A) The intermittent control model. (B) The forced-choice continuous-like control model. (A) Characteristics of the unbiased synaptic weight vector *W*_unbiased_: When the pendulum is in S_1_ as (A)-1, the backward and forward population of Go-striatum receives strong and weak inputs, respectively, and the backward and forward population of NoGo-striatum receives weak and strong inputs, respectively. When the pendulum is in S_3_, as (A)-3 the forward population of the Go-striatum and the backward population of the NoGo-striatum receives strong inputs and the backward population of the Go-striatum, and the forward population of the NoGo-striatum receives weak inputs. When the pendulum is in S_2_ or S_4_ as (A)-2 and (A)-4, the sensory cortico-striatal inputs to the antagonistic populations were set with equal strength. (B) Characteristics of the biased synaptic weight vector *W*_biased_: When the pendulum is in S_1_ or in S_4_ as (B)-1 and (B)-4, the backward and forward population of Go-striatum receives strong and weak inputs, respectively, and the backward and forward population of NoGo-striatum receives weak and strong inputs, respectively. When the pendulum is in S_2_ or S_3_ as (B)-2 and (B)-3, the forward population of the Go-striatum and the backward population of the NoGo-striatum receives strong inputs and the backward population of the Go-striatum, and the forward population of the NoGo-striatum receives weak inputs.

The exploration of the sensoristriatal synaptic weight vector *W*_unbiased_ to implement the intermittent control model was performed under the small fixed feedback parameters to (*P,D*) = (0.45*mgh*, 18), with which the upright equilibrium remains unstable (unstable focus) even if the underlying PD controller operates continuously. Moreover, the noise amplitude was set to *σ* = 0.2 Nm. These values were consistent with the feedback gains and noise intensity identified by the data assimilation of postural sway data from healthy people (Suzuki et al. 2020).

#### Forced-Choice (Continuous-Like) controller --Baseline Regime

To investigate the neural origin of sway-related beta modulation, we compared the balanced intermittent control regime (*W*_unbiased_) against a sustained active control condition, termed the forced-choice (continuous-like) regime, in which the sensoristriatal synapse leads to biased action selections by the use of biased synaptic weight vector, referred to as *W*_biased_.

It is important to clarify that this forced-choice regime is implemented within the identical spiking CBGT network structure by biasing the corticostriatal weight matrix (*W*_biased_) toward the Direct (Go) pathway. This configuration forces the cortical population to persistently cross the firing-rate threshold and generate active torque whenever the body deviates from equilibrium, thereby eliminating the prolonged Go/NoGo competitive dynamics (eliminating or narrowing DT) seen in safe sway regions (*S*_2_, *S*_4_). This regime serves as an internal computational baseline to evaluate how LFP spectral properties change when subcortical decision time is minimized, rather than representing a literal implementation or critique of classical continuous feedback control models, e.g., (Peterka 2002).

The forced-choice controller was implemented by exploring *W*_biased_ that enforces a biased choice across all regions of the phase plane, thereby minimizing the null-action periods. When the pendulum is in S_1_ (tilted forward, falling forward) or in S_4_ (tilted forward, moving backward), the network parameters favor backward torque by routing strong inputs to the backward Go-striatum and forward NoGo-striatum, and weak inputs to their counterparts (Fig. 3B-1, Fig. 3B-4; Table 3B). Conversely, in regions S_2_ (tilted backward, moving forward) and in S_3_ (tilted backward, falling backward), the projections favor forward torque (Fig. 3B-2, Fig. 3B-3; Table 3B). Under this parameter configuration, cortical action selection resembles a forced choice between opposing active torques. Consequently, the DT is markedly shortened, the probability of selecting the null action becomes negligible, and the system operates as a continuous controller.

To prevent delay-induced instability inherent to continuous feedback architectures, the feedback gains must be substantially higher than those of the intermittent controller (Asai et al. 2009). We therefore used (*P,D*) = (0.5*mgh*, 150), which yields a stable upright equilibrium under uninterrupted feedback. Moreover, the noise amplitude was increased to *σ* = 0.6 Nm. These values are in accordance with the previous continuous control simulations to reproduce postural sway in patients with Parkinson’s disease (Suzuki et al. 2020).

#### Physiological and Computational Rationales for System Parameters

The key operational parameters of the CBGT closed-loop model were defined based on established neurophysiological and biomechanical empirical data:

- **Decision Threshold for Torque Generation:** Action selection was modeled using a firing-rate accumulator threshold mechanism (Tanaka 2007). Active torque generation was triggered when M1 population firing rates crossed a decision threshold. While set to 50 Hz in main simulations, sensitivity analyses confirmed that lowering the threshold to 40 Hz produced identical, qualitative sway-phase-locked beta-band ERD/ERS modulations, confirming that beta dynamics are robust and independent of specific threshold values.
- **Temporal Smoothing and Delays:** Cortical output firing rates were calculated using a 0.1-s moving average window to model the postsynaptic integration kinetics of cortical and motoneuronal dendrites. A total loop latency of 200 ms was incorporated to represent the long-loop transcortical/CBGT feedback delay encompassing sensory processing, subcortical action selection, and neuromuscular lag (Loram et al. 2005b).
- **Biomechanical Constraints:** Passive ankle joint stiffness was set to 80% of gravitational overturning torque (*mgh*), reflecting empirical measurements in humans showing that passive tissue elasticity alone is insufficient for upright stability (Loram and Lakie 2002; Casadio et al. 2005). Sway dynamics 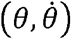 were evaluated relative to the upright equilibrium state (*θ* = 0) (Peterka 2002). Sensitivity analyses confirmed that that varying passive stiffness within the physiologically reported range, 80% to 90% of *mgh* (Loram and Lakie 2002), produced consistent postural stabilization and robust, identical sway-phase-locked beta ERD/ERS dynamics.

### Simulation protocols and data analysis

#### Simulation parameters and testing trials

We defined a localized rectangular boundary in the phase plane (0 ≤ |*θ*| ≤ 0.05 rad and 0 ≤ |*ω*| ≤ 0.05 rad/s) centered at the origin; trajectories escaping this domain were classified as falls (postural failure). Each trial was initiated from a random position within a narrow region (0≤ |*θ*| ≤ 0.02 rad and 0≤ |*ω*| ≤ 0.02 rad/s). A simulation trial was considered successful if the pendulum avoided falling for 205 s, and this protocol was repeated to gather 30 successful trials. Data from the first 5 s of each trial were excluded from analysis to eliminate transient initialization artifacts.

#### Simulated EEG reconstruction

To assess beta-band desynchronization and synchronization, we simulated cortical EEG using the LFP generated by the motor cortex. The LFP signal was computed by summing all excitatory and inhibitory synaptic currents received across the entire cortical population (Ouyang et al. 2022):

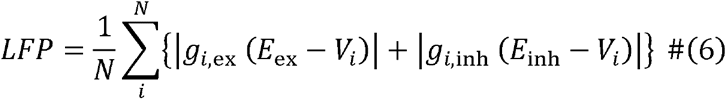

where *g*_i,ex_, *g*_i,inh_, and *v*_i_ represent the time-dependent excitatory synapse conductance, the time-dependent inhibitory synapse conductance, and the membrane potential (served as a dynamic variable) of th cortical neuron, respectively. (*E*_ex_ and *E*_inh_ are constants as defined above.) Neurons from both the backward and forward cortical sub-populations were pooled indiscriminately, yields a total population size of *N* = 1,000.

#### Sensitivity Analysis of Corticostriatal Weights and Decoupling from Mechanical Gains

To evaluate the sensitivity and robustness of the intermittent control regime and sway-related beta modulation to corticostriatal projection strengths, we performed a systematic parameter sweep. The key functional difference between the unbiased condition (*W*_unbiased_) and the biased/forced-choice condition (*W*_biased_) lies in the eight-dimensional weight vector *W* governing sensory cortical inputs to striatal Direct (Go) and Indirect (NoGo) pathways across phase regions (*S*_2_ and *S*_4_).

We defined a linear interpolation path in this 8D weight space parameterized by *s* ∈ [0,1] :

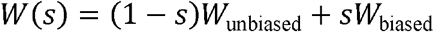

where *s* = 0 corresponds to *W*_unbiased_ and *s*= 1 to *W*_biased_. We evaluated five evenly spaced intermediate weight vectors (*W*_1_ to *W*_5_ corresponding to *s* = 1/6,2/6,3/6,4/6,5/6, respectively).

To test whether beta modulation depends solely on synaptic weight design or on the closed-loop interaction between neural competition and physical body movement, we independently fully crossed these seven weight configurations (*W*_unbiased_, *W*_1_--*W*_5_,*W*_biased_) with two distinct mechanical feedback parameter sets (*P,D, σ*):

1. **Healthy/Small feedback regime (*P***_small_,*D*_small_, *σ*_small_):, (*P,D*) = 0.45*mgh*,18), noise amplitude *σ*= 0.2 Nm/rad, derived from data assimilation of healthy human sway during quiet standing (Suzuki et al. 2020).
2. **PD/Large feedback regime *p***_large_, *D*_large_, *σ*_large_) :, (*P,D*) = (0.5*mgh*,150), noise amplitude *σ*= 0.6 Nm/rad, representing the hyper-stiffness and exaggerated noise observed in severe Parkinsonian postural impairment (Suzuki et al. 2020).

#### Evaluation of the CBGT circuit topology (Experiment 2)

In *Experiment* 1, the network architecture could not be altered because the CBGT controller was strictly constrained by the requirement to stabilize the pendulum. To systematically evaluate how specific loops within the CBGT circuit shape the simulated EEG, *Experiment 2* introduced targeted structural dissections and parameter modifications. To maintain consistent sensory drive despite the loss of postural stability caused by these perturbations, the pendulum was driven passively along a predefined periodic trajectory, decoupled from the cortical motor command.

The sensory feedback pathway remained intact, allowing state-dependent signals from the sensory cortex to drive the network exactly as in *Experiment 1*. For conditions utilizing the intermittent CBGT parameters, i.e., with the unbiased synaptic weight vector *W*_unbiased_, i.e., Conditions 1 (the intermittent CBGT controller as is), 3 (firing rate of sensory cortex is altered to λ=200), 4 (projections from the sensory cortex to the motor cortex and the striatum were dissected), 5 (projections from the motor cortex to STN were dissected), and 6 (the local feedback projections from GPe to STN were dissected), the passive trajectory alternated between regions S_1_ and S_4_ with a 5-second period (switching every 2.5 s), mimicking the characteristic phase-plane trajectories observed in *Experiment 1* (Fig. 6A). For the forced-choice continuous-like control configuration, i.e., with the biased synaptic weight vector *W*_biased_, (Condition 2), the passive trajectory routed the system through S_1_ and S_4_ for 1.25 s each, followed by S_3_ for the remaining 2.5 s (total period of 5 s), approximating the elliptical trajectory in Fig. 6B. Each simulation ran for 5,000 s to yield 1,000 complete cycles (epochs) for robust spectral analysis. We evaluated the following six distinct network conditions:

**[Condition 1]** Baseline intermittent CBGT controller with *W*_unbiased_ under passive periodic movement between S_1_ and S_4_ (period: 5 s).

**[Condition 2]** Baseline forced choice continuous-like CBGT controller with *W*_biased_ under passive periodic movement through S_1_, S_4_ and S_3_ (period: 5 s).

**[Condition 3]** Same configuration as Condition 1 with *W*_unbiased_, but with the sensory cortex firing rate increased to *λ*=200 Hz to analyze the impact of sensory signal intensity.

**[Condition 4]** Same configuration as Condition 1 with *W*_unbiased_, but with the thalamo-cortical projections disconnected to open the major cortico-basal ganglia feedback loop.

**[Condition 5]** Same configuration as Condition 1 with *W*_unbiased_, but with the pathway from the cortex to the STN disconnected, eliminating STN activity to simulate a network lacking the hyperdirect pathway.

**[Condition 6]** Same configuration as Condition 1 with *W*_unbiased_, but with the local inhibitory connections from the GPe and the STN disconnected (no GPe-STN loop).

#### Signal processing and statistical averaging

Time-series data—including pendulum kinematics, active torques, and cortical LFPs—were extracted over cumulative durations of 2,050 s for *Experiment 1* and 5,050 s for *Experiment 2* (combining 10 separate simulation runs per condition, discarding the initial 5 s transient phase).

For *Experiment 1*, individual fall-recovery cycles were isolated. Each cycle was defined as beginning at the onset of a micro-fall and terminating at the completion of the subsequent micro-recovery. To isolate prominent postural adjustments, the start and end points of these cycles were detected using the scipy.signal.find_peaks() function in Python, applied to the absolute state vector |*θ* (*t*) − *ε*| where *ε*=0.001 rad is a small baseline offset used to focus trajectory-arcs locating slightly away from the upright position. We extracted cycles exhibiting a sway amplitude greater than 0.001 rad and a duration longer than 0.05 s to filter out minor chattering. For *Experiment 2*, epoch segmentation was defined directly by the fixed 5-second period of the passive driving trajectory.

Time-frequency representations of the LFP were computed using a short-time Fourier transform (STFT) with a 1-second sliding window. To analyze event-related spectral perturbations (ERSPs), the time-averaged power across the entire trial was established as the baseline for each frequency, and dynamic power fluctuations were converted to decibels (dB) relative to this baseline (Nakamura et al. 2023). Because individual fall-recovery cycles in *Experiment 1* varied in duration, the kinematics, torque profiles, and ERSP matrices were time-warped. The micro-fall and micro-recovery intervals were normalized separately so that their lengths matched the respective mean ensemble durations across all extracted epochs before computing the final event-locked average. For *Experiment 2*, ERSPs and population-specific firing rates were directly ensemble-averaged over the fixed 1,000 epochs. All computational modeling and data analysis pipelines were executed using Python (version 3.10.13).

## Results

### Distinct postural stabilization and sway kinematics driven by intermittent and continuous CBGT Controllers

To evaluate the behavioral consequences of different neural control strategies, we first analyzed the phase-plane trajectories and power spectral densities (PSDs) of the inverted pendulum stabilized by the intermittent (with *W*_unbiased_) and forced-choice continuous-like (with *W*_biased_) CBGT controllers (Fig. 4). The intermittent control model exhibited a characteristic butterfly-wing-shaped trajectory in the phase portrait (Fig. 4A, upper panel). This pattern indicates that the pendulum achieved stability by exploiting the stable manifold of the saddle-type upright equilibrium point, as evidenced by the hyperbolic arcs centered at the origin during the control-off (decision time, DT) periods (represented by black segments). Furthermore, the low-frequency band of the angle PSD for the intermittent model exhibited a 1/f-like power-law scaling (Fig. 4A, lower panel), a well-established computational hallmark of intermittent control strategies (Asai et al. 2009; Nomura et al. 2022).

**Figure 4:**
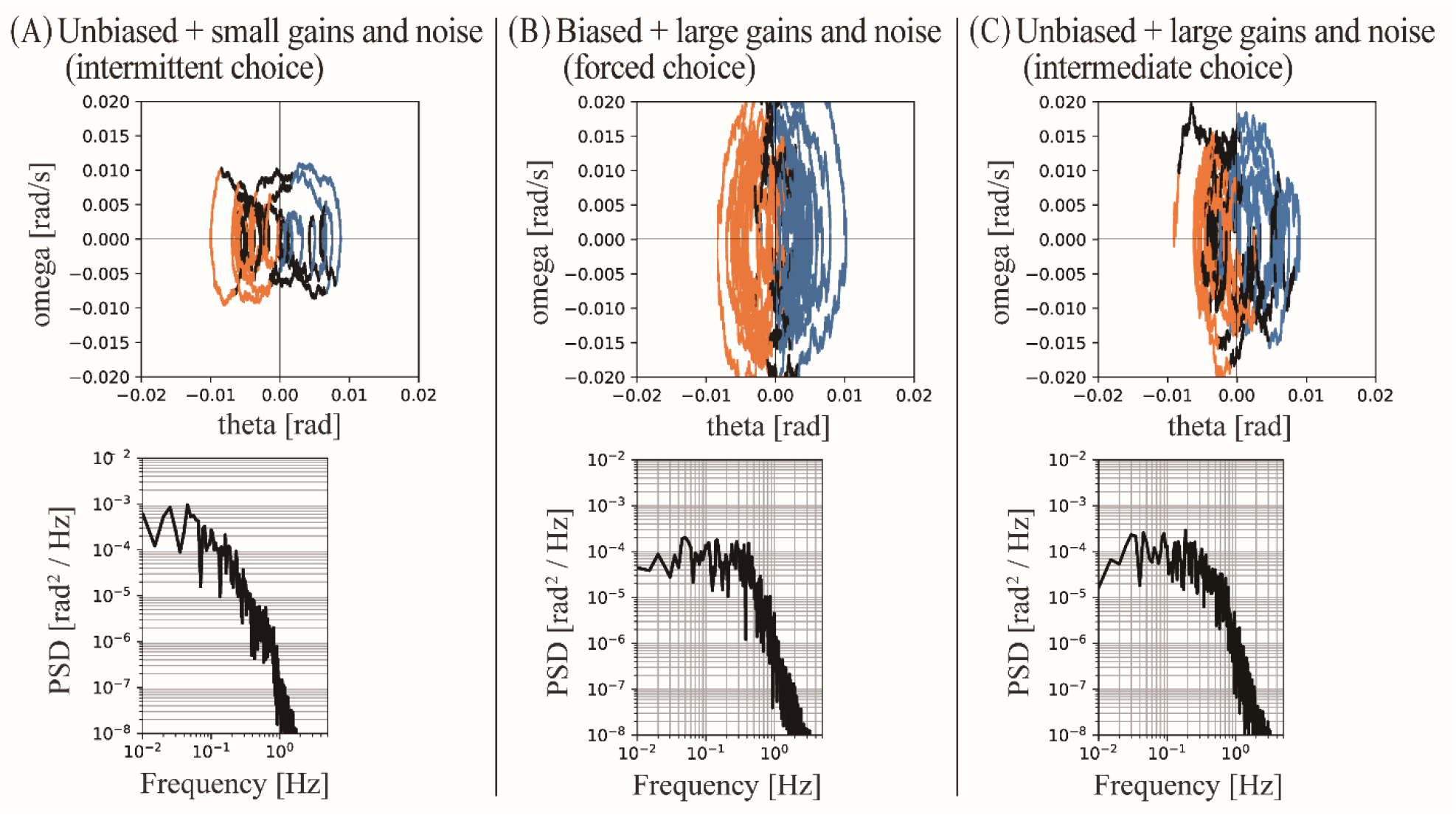
Dynamics of the inverted pendulum controlled by variants of the CBGT controllers. (A) The intermittent control model implemented with unbiased weight vector *W*_unbiased_ and small feedback gains and noise (*P*_small_, *D*_small_, *σ*_small_). (B) The forced-choice continuous-like control model implemented with biased weight vector *W*_biased_ and large feedback gains and noise (*P*_large_, *D*_large_, *σ*_large_). (C) A type of intermediate-choice model with unbiased weight vector *W*_unbiased_ and large feedback gains and noise (*P*_large_, *D*_large_, *σ*_large_). The upper panels are the phase portrait with a simulated trajectory for 40 seconds. Blue, orange and black segments represent dynamics of the pendulum controlled by the backward-torque action, the forward-torque action, and the null action of the CBGT controller, respectively. Lower panels are ensemble averaged PSD of the tilt-angle time series for 200 seconds across 30 simulation trials.

In contrast, the forced-choice continuous-like control model displayed an elliptical spiral trajectory (Fig. 4B, upper panel). In this regime, the system dynamics were continuously dominated by alternating active commands—backward torque in the right half-plane and forward torque in the left half-plane—with negligible null-action intervals between them (Fig. 4B, upper panel). Reflecting this uninterrupted feedback regulation, the low-frequency regime of the angle PSD exhibited a plateau characteristic of white noise (Fig. 4B, lower panel). This profile closely mimics the atypical, rigid postural sway patterns documented in patients with advanced Parkinson’s disease (Suzuki et al. 2020; Nomura et al. 2022). Together, these behavioral simulations confirm that functionally tuning the cortico-striatal synaptic weights is sufficient to switch the closed-loop system between a biomimetic intermittent control state and a pathological continuous stiffness state.

### Sway-coupled spiking modulations and phasic cortical LFPs emerged solely under intermittent control (Experiment 1)

To uncover the subcortical circuit mechanisms driving sway-locked beta modulations, we examined the model-generated internal neural dynamics of the CBGT network (Fig. 5). It should be emphasized that while body sway kinematics 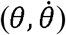 and scalp EEG beta modulations are directly validated against empirical human postural data, the subcortical population firing rates and striatal LFP patterns shown in Figures 5 and 6 represent computational model predictions that provide testable hypotheses for future neurophysiological investigations.

**Figure 5:**
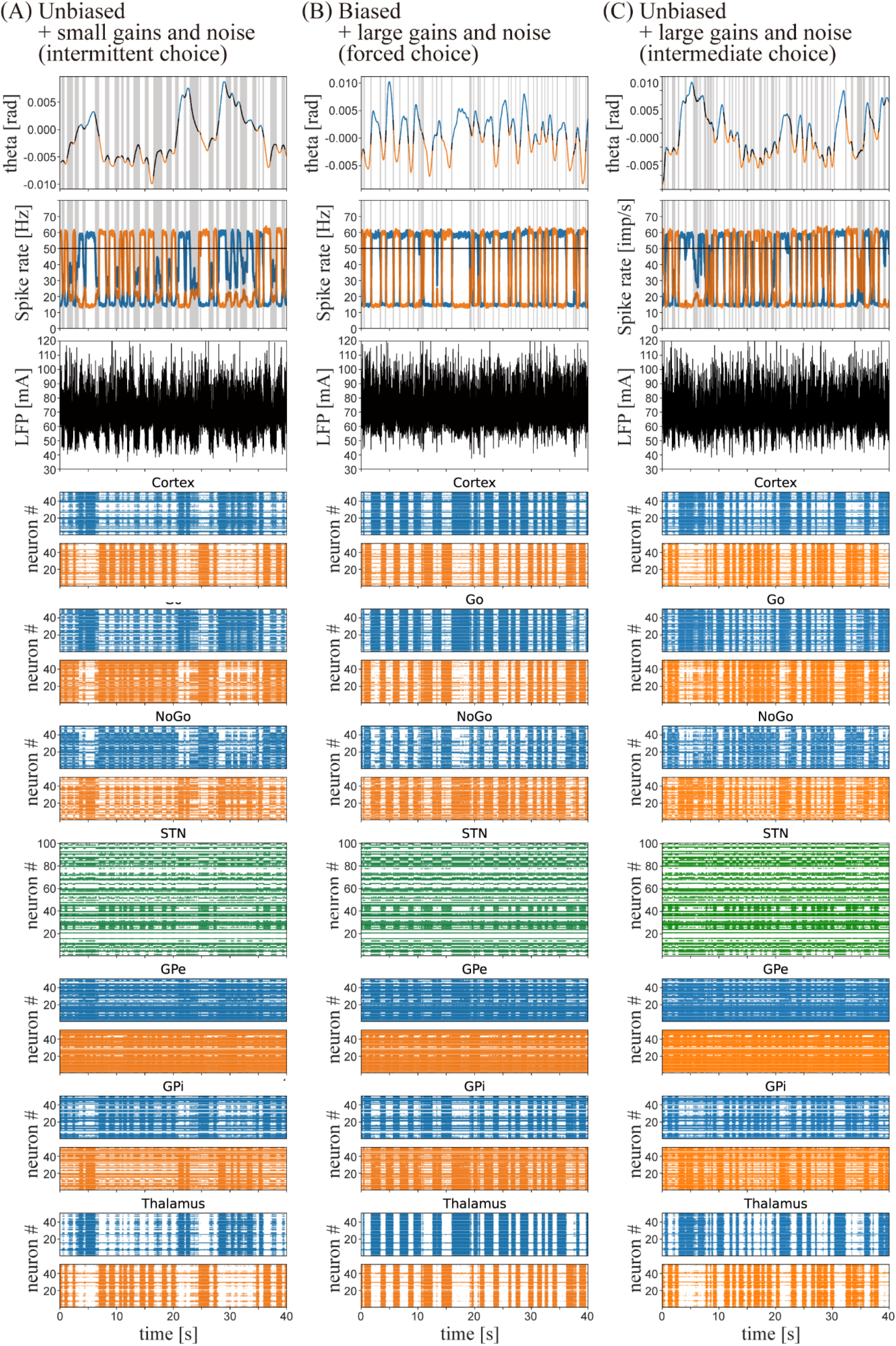
Model-simulated sway and internal neural dynamics over multiple fall-recovery cycles. (A) The intermittent control model implemented with *W*_unbiased_ and (*P*_small_, *D*_small_, *σ*_small_). (B)The forced-choice continuous-like control model implemented with *W*_biased_ and (*P*_large_, *D*_large_, *σ*_large_). (C) A type of intermediate-choice model with w_unbiased_ and (*P*_large_, *D*_large_, *σ*_large_). Top trace: The time series of the tilt angle with the color-codes representing the choice of actions as in Fig. 4. Second trace: Spike rates of the backward and forward populations in the cortex. Gray-shaded vertical bars represent a sequence of decision times (DT). Third trace: LFP as the simulated EEG. Panels at the 4th to the 10th row (expect 7th). Raster plots of spikes for 50 neurons out of 500 neurons in each of the backward and forward populations for the cortex, Go-striatum, NoGo-striatum, STN, GPe, GPi and thalamus. The 7^th^ row is for 100 STN neurons out of 1,000 neurons.

**Figure 6:**
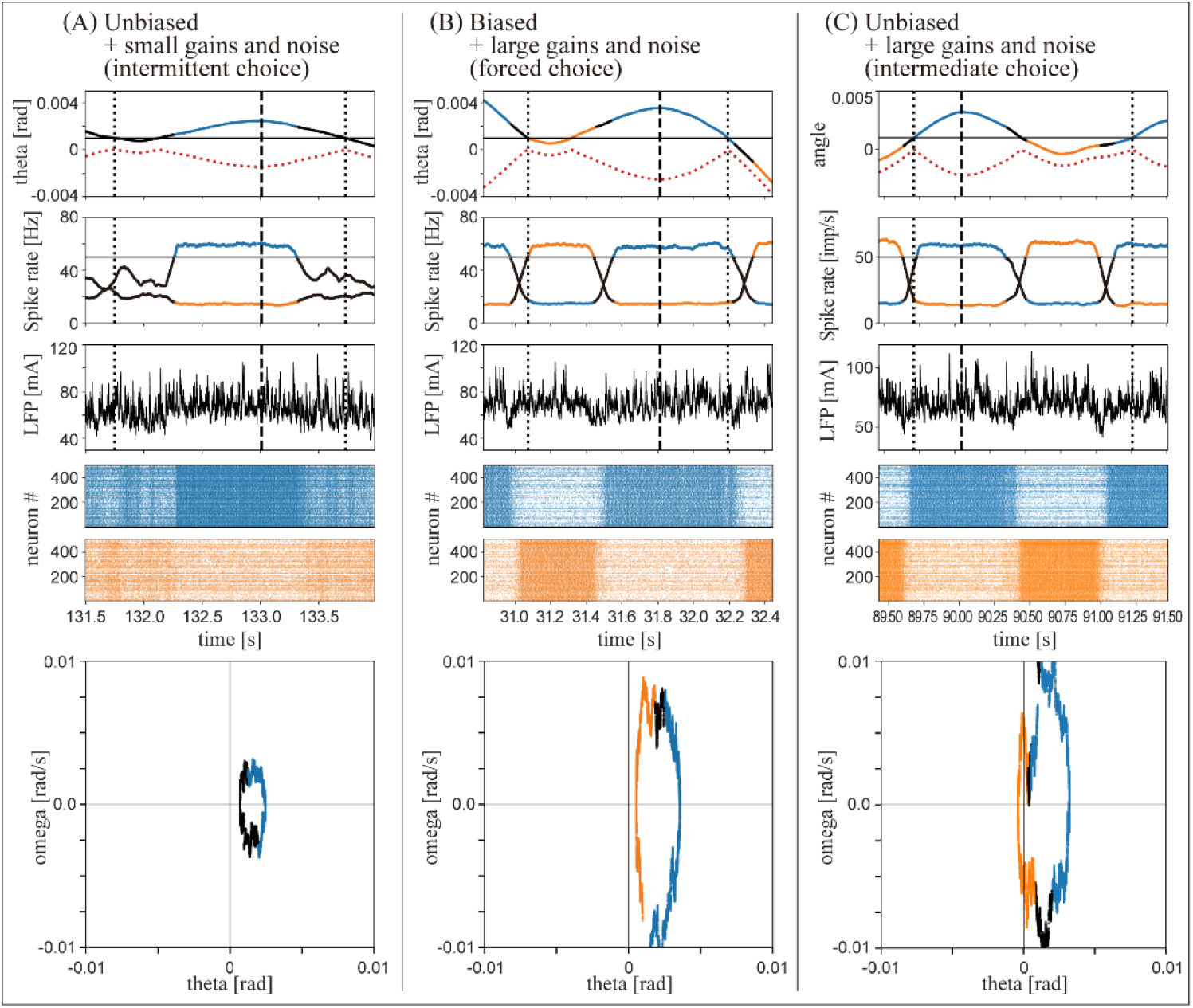
Model-simulated sway and neural dynamics for a single extracted fall-recovery cycle along with the accompanying changes in the spike rates. (A) The intermittent control model with *W*_unbiased_ and (*P*_small_, *D*_small_, *σ*_small_). (B) The forced-choice continuous-like control model with *W*_biased_ and (*P*_large_, *D*_large_, *σ*_large_). (C) A type of intermediate-choice model with *W*_unbiased_ and (*P*_large_, *D*_large_, *σ*_large_). Spike rates of the backward and forward cortical populations (2^nd^ row), and the corresponding LFP time series (3^rd^ row) and raster plots of cortical neurons (4^th^ row), along side with an extracted tilt angle profile (1^st^ row). The phase portraits in the bottom row illustrate the trajectories of these individual fall-recovery cycles. Color codes for segments represent the three choices of actions as in Figs. 4 and 5. Dotted curves below the tilt angle profile represent the the time series data of absolute value |*θ* (*t*)− *ε*| where *ε*=0.001 (the offset) used for the cycle extraction.

In both intermittent and forced-choice continuous-like control models, the alternation between forward and backward active torques was closely coupled with reciprocal firing rate modulations within the motor cortex (Fig. 5A-B, top and second rows). Specifically, active torques were executed when the spike rates of the corresponding cortical populations reached the 50 Hz decision threshold. However, a fundamental difference emerged in the temporal structure of these neural commands. The intermittent control model displayed clear, prolonged intervals during which neither the forward nor backward cortical population reached the threshold (Fig. 5A, top and second rows). These gaps successfully defined the control-off decision time (DT). In contrast, the forced-choice continuous-like control model exhibited immediate, antagonistic transitions: as soon as one population dropped below the threshold, the other crossed it without delay (Fig. 5B, top and second rows).

This difference in network operation profoundly impacted the simulated local field potentials (LFPs) in the motor cortex (Fig. 5A-B, third row). The intermittent model generated robust, phasic LFP modulations locked to the postural sway cycle. Conversely, the forced-choice continuous-like model generated tonic, unmodulated LFP activity, where the amplitude of sway-coupled modulation was markedly attenuated (Fig. 5, right). This loss of phasic complexity was further reflected in the population-wide raster plots (Fig. 5A-B, lower panels). While the intermittent network sustained high-complexity, structured firing patterns across all nuclei, the forced-choice network fell into stereotyped, simple alterations between antagonistic populations.

To better capture these dynamics at a single-action resolution, we isolated individual fall-recovery cycles (Fig. 6). In the intermittent model, each cycle began with a hyperbolic segment near the upright position during the control-off DT phase (black), followed by a clockwise arc driven by a discrete burst of backward torque (blue) (Fig. 6A, phase portrait). This sequence was supported by well-timed, transient escalations in cortical firing rates and discrete LFP bursts (Fig. 6A, temporal traces). Conversely, the fall-recovery cycle in the forced-choice model consisted of transitions directly from forward-torque (orange) to backward-torque actions (blue) via a negligible null-action gap (Fig. 6B, phase portrait). This confirms that forced-choice continuous-like control flattens the rich temporal dynamics of the CBGT network into rigid, tonic activity.

### Post-movement beta rebound (ERS) is exclusively regenerated by the intermittent Control loop (Experiment 1 continued)

To determine whether our embodied CBGT network replicates human scalp EEG signatures, we analyzed event-related spectral perturbations (ERSPs) computed from the motor cortex LFPs during the isolated fall-recovery cycles (Fig. 7). The dynamic modulation of the beta band (13–25 Hz) provided the pivotal finding of this study (Fig. 7, upper row).

**Figure 7:**
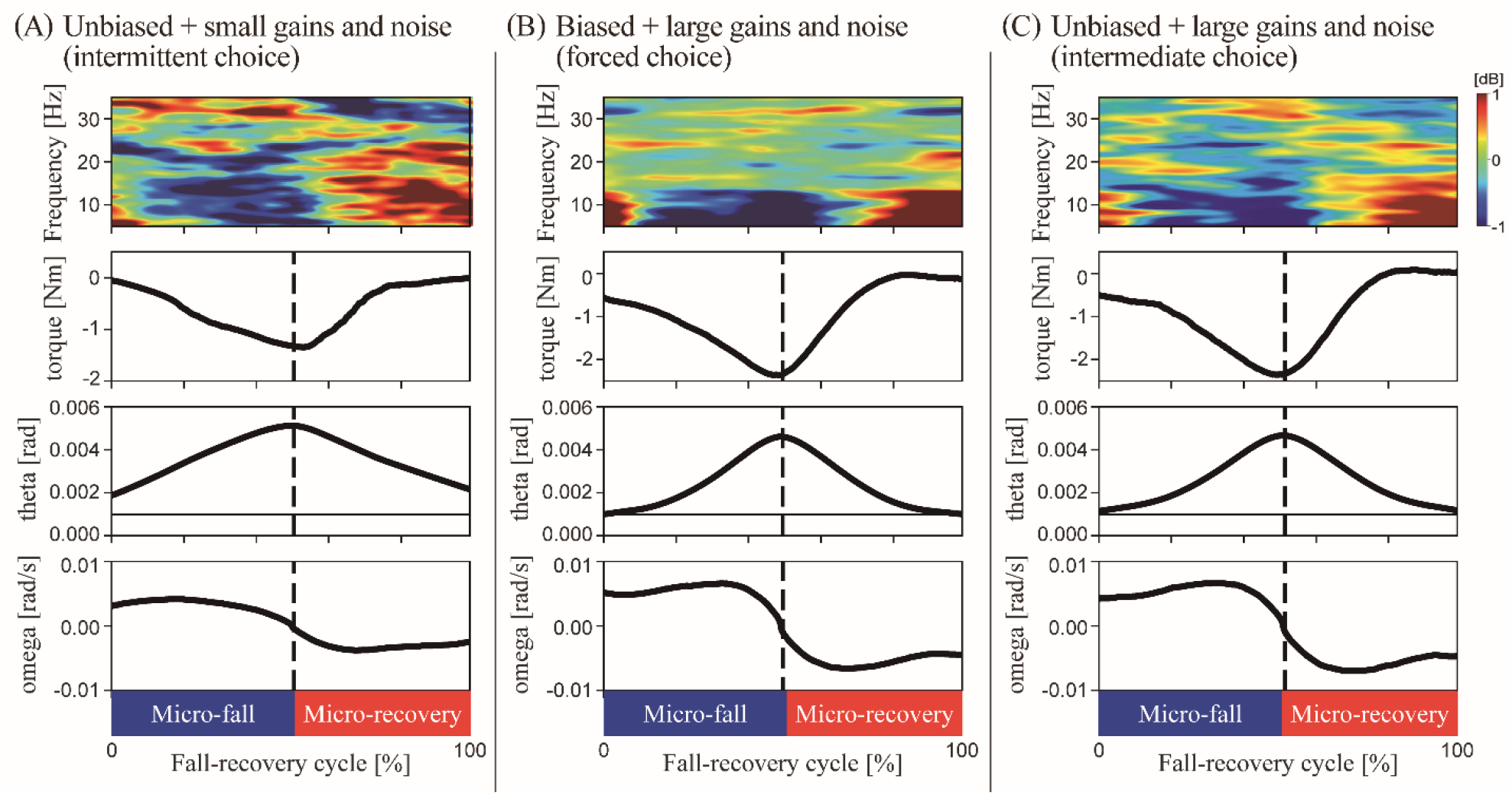
Beta ERSP locked to the micro-fall/recovery cycle for the intermittent CBGT controller, but no beta modulation for the forced-choice CBGT controller. (A) The intermittent control model with *W*_unbiased_ and (*P*_small_, *D*_small_, *σ*_small_). (B) The forced-choice continuous-like control model with *W*_biased_ and (*P*_large_, *D*_large_, *σ*_large_). (C) A type of intermediate-choice model with *W*_unbiased_ and _large_, *D*_large_, *σ*_large_). ERSP of simulated EEG and dynamics of the pendulum averaged across all fall-recovery cycles for each control model in Experiment 1. Only forward falling cycles were used for the ensemble average. Top: Ensemble averaged ERSPs of reconstructed LFPs over all fall-recovery cycles. 2^nd^ row: Control torque applied to the pendulum. 3^rd^ and 4^th^ rows: angle and angular velocity of the pendulum, respectively. 0% and 100% in horizontal axis of for all panels represent the onset of the micro-fall and offset of the micro-recovery, respectively. Vertical dashed lines represent the division between the micro-fall and the micro-recovery.

In the intermittent control model, a prominent beta event-related desynchronization (beta-ERD) occurred during the forward micro-fall phase, coinciding with the cortical preparation and execution of backward torque to arrest the pendulum (Fig. 7A-top). Crucially, during the subsequent micro-recovery phase—after the pendulum’s velocity reversed and the active torque returned to zero (the control-off DT window)—a striking beta event-related synchronization (beta-ERS, or post-movement beta rebound) emerged (Fig. 7A-top).

In stark contrast, this beta-band rhythmicity was completely abolished in the forced-choice control model (Fig. 7B-top). The continuous architecture failed to exhibit any statistically significant beta-ERD or beta-ERS across the normalized sway cycle. Interestingly, both models successfully replicated power modulations in the lower frequency bands (theta and alpha, 5–13 Hz), showing ERD during micro-falls and ERS during micro-recoveries. In the high-frequency range (30–35 Hz), both models also captured micro-fall ERS and micro-recovery ERD, though it was less pronounced in the continuous regime. These spectral profiles demonstrate that cortical beta modulations—specifically the post-movement beta rebound—do not merely reflect passive sensory registration, but are a distinct neurocomputational signature of intermittent motor selection.

### Phasic neuronal population modulations and spike-rate profiles across the CBGT network (Experiment 1 continued)

To elucidate the mechanisms underlying the generation of beta-ERD and beta-ERS observed in the ERSP of the simulated EEG (LFP) shown in Fig. 7, we tracked the ensemble firing rates of the backward and forward populations within each CBGT nucleus across the fall-recovery cycle (Fig. 8).

**Figure 8:**
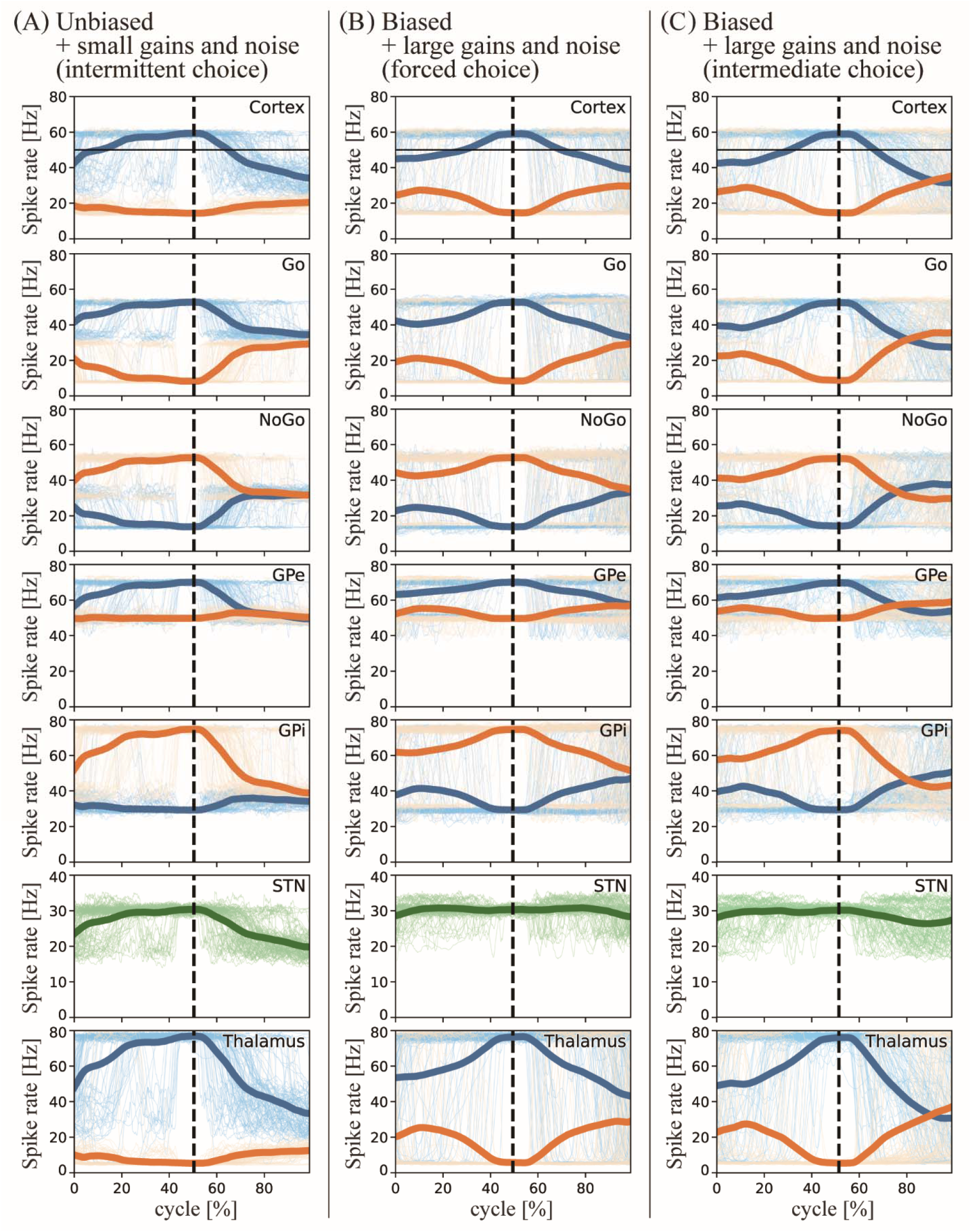
Spike rate dynamics in the CBGT controllers. (A) The intermittent control model with *W*_unbiased_ and (*P*_small_, *D*_small_, *σ*_small_). (B) The forced-choice continuous-like control model with *W*_biased_ and (*P*_large_, *D*_large_, *σ*_large_). (C) A type of intermediate-choice model with *W*_unbiased_ and (*P*_large_, *D*_large_, *σ*_large_). Spike rate dynamics of the backward (blue) and forward (orange) populations in the cortex, Go-striatum, NoGo-striatum, GPe, GPi, STN and thalamus in Experiment 1. In each panel, Thin curves represent the spike rates of 100 single cycles and thick curves represent the spike rates of ensemble average of all cycles.

In the intermittent control model (Fig. 8A), the temporal evolution of cortical spike rates was closely coupled with the postural sway cycle. During the forward micro-fall phase, the spike rate of the backward-action population was elevated, remaining near or above the 50 Hz decision threshold, whereas it decreased to approximately 30–35 Hz during the subsequent micro-recovery phase. Conversely, the spike rate of the forward-action population remained low at approximately 20 Hz throughout the entire fall-recovery cycle.

The cortical local field potential (LFP) is defined as the summation of postsynaptic currents (PSCs) across all cortical neurons. These PSCs integrate inputs from the thalamocortical projections, a continuous 100-Hz sensory feedback signal, and reciprocal inhibition between the two antagonistic cortical populations. Importantly, the frequency components of the LFP strongly reflect the underlying spike rates of all cortical and thalamic neurons.

These cortical spike rates were heavily influenced by excitatory projections from the thalamus; indeed, the spike-rate modulations of the two thalamic populations were qualitatively identical to those observed in the cortex. When evaluating the frequency components of the LFP based on the mean spike rates of both the backward and forward populations across the cortex and thalamus, the aggregate spike rate averaged approximately 35–40 Hz during the micro-fall phase and decreased to 20–25 Hz during the micro-recovery phase. This attenuation in the aggregate firing rate indicates that power within the beta band of the cortical LFP increases during the micro-recovery phase, thereby generating the post-movement beta ERS.

Furthermore, the spike rates of the two populations in the GPi, which provides inhibitory output to the thalamus, exhibited modulations that mirrored the thalamocortical activity, albeit with the backward and forward populations reversed. Specifically, during the micro-fall phase, the spike rate of the forward population in the GPi was high and subsequently decreased during the micro-recovery phase, while the backward GPi population maintained a nearly constant low spike rate throughout the cycle. The spike-rate changes in the two GPe populations, which inhibit the GPi, mirrored the modulations of the GPi.

A prominent characteristic shared across the cortex, thalamus, GPi, and GPe was that the spike rates of individual populations, as well as the aggregate firing rates of both populations within each nucleus, exhibited clear phasic modulations coupled with the fall-recovery cycle. These phasic dynamics appear to be driven by the STN, which provides excitatory projections to both the GPe and GPi and exhibits robust phasic spike-rate modulations (see the results for the STN-dissected experiments described below in Figs. 10 and 11).

In contrast, in the forced-choice control model (Fig. 8B), the cortical spike rates followed a distinct and less modulated temporal profile. During the forward micro-fall phase, the backward population maintained a persistently high spike rate near the 50 Hz decision threshold, peaking at approximately 60 Hz near the transition from the fall to the recovery phase, and remained similarly elevated during the subsequent micro-recovery phase. Meanwhile, the spike rate of the forward population mirrored the temporal profile of the backward population around a horizontal baseline of approximately 35 Hz. Consequently, the mean aggregate spike rate of the combined backward and forward cortical populations remained nearly constant at approximately 35 Hz throughout the entire fall-recovery cycle. Because the spike-rate modulations of the two thalamic populations were qualitatively identical to those of the cortex, no beta-band power modulation occurred.

The spike-rate profiles in all other nuclei of the forced-choice control model were qualitatively similar to those observed in the cortex. Notably, the STN spike rate remained nearly constant throughout the fall-recovery cycle, exhibiting enhanced tonic spiking activity fixed at approximately 30 Hz.

### Decoupling synaptic weights and mechanical dynamics: Beta modulation as an emergent closed-loop property

To address whether the observed sway-related beta modulation is an artifact manually hard-coded via the corticostriatal weight matrix (*W*_unbiased_) or an emergent property of closed-loop body-brain dynamics, we conducted systematic sensitivity analyses and decoupled the neural synaptic weights from the mechanical feedback gains (Figs 4C–8C and Fig. 9).

**Figure 9:**
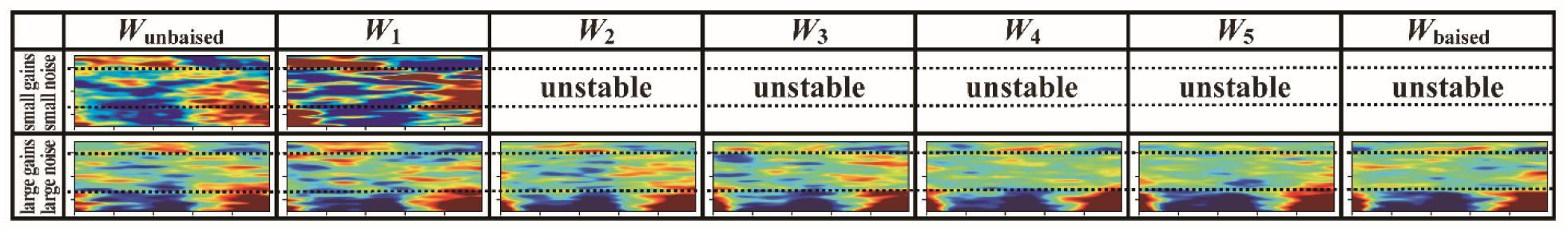
Sensitivity analysis of corticostriatal weights and mechanical gains. ERSP was examined for varied corticostriatal weight vector *W* ranging between *W*_unbiased_ and *W*_biased_ across five equally spaced intermediate points *W*_1_ (close to *W*_unbiased_) to *W*_5_ (close to *W*_biased_). Top row: Under the flexible mechanical regime with small gains and noise (*P*_small_, *D*_small_, *σ*_small_). Bottom row: Under the rigid mechanical regime with large gains and noise (*P*_large_, *D*_large_, *σ*_large_). Stability was achieved *exclusively* with *W*_unbiased_ and *W*_1_ for (*P*_small_, *D*_small_, σ_small_). For (*P*_large_, *D*_large_, *σ*_large_), sway-related beta modulation vanished for all examined weight vectors, even when *W*_unbiased_ was preserved but combined with large mechanical gains and noise (*P*_large_, *D*_large_, *σ*_large_).

### Postural Stability as a Criterion for Weight Selection

We first interpolated the eight-dimensional corticostriatal weight vector *W* between *W*_unbiased_ (*s*= 0) and *W*_biased_ (*s*= 1) across five intermediate points (*W*_1_ to *W*_5_; *s* = 1/6,…,5/6). Under the healthy mechanical regime with small gains and noise (*P*_small_, *D*_small_, *σ*_small_), successful stabilization of the inverted pendulum was achieved *exclusively* with *W*_unbiased_ and *W*_1_ (Fig. 9). All other weight configurations (*W*_2_ through *W*_5_ and *W*_biased_) failed to maintain upright stance, leading to mechanical falling (Fig. 9). This demonstrates that *W*_unbiased_ was constrained *a priori* by the functional requirement of postural balance, rather than tuned *a posteriori* to reproduce beta modulation.

### Quenching of Beta Modulation Under High Sway Velocity

Crucially, when *W*_unbiased_ was preserved but combined with large mechanical gains and noise (*P*_large_, *D*_large_, *σ*_large_), sway-related beta modulation completely vanished (Fig. 7C), and the system yielded dynamics virtually indistinguishable from the forced-choice model (Figs. 4C–8C and Fig. 9). Although *W*_unbiased_ provided balanced Go/NoGo inputs in safe phase regions (*S*_2_, *S*_4_), the high angular velocity driven by large *P*-*D* gains forced the body’s sway state to rapidly pass through *S*_2_ and *S*_4_. Consequently, cortical firing rates rapidly crossed the 50-Hz threshold, depriving the CBGT circuit of sufficient physical time to maintain prolonged decision time (DT) (Figs. 5C, 6C, and 8C). Without sustained Go/NoGo competitive dynamics, the circuit failed to generate the LFP spectral fluctuations required for beta ERD/ERS (Fig. 7C).

These results confirm that sway-related beta modulation is not a built-in consequence of *W*_unbiased_ alone, but an emergent signature produced only when balanced CBGT attractor dynamics and moderate physical sway velocities are mutually matched.

### Structural dissection and parameter manipulations reveal closed-loop mechanisms underpinning LFP frequency modulations (Experiment 2)

In Experiment 2, to elucidate the network mechanisms underlying the sway-related beta modulations (ERD and ERS) observed under the intermittent control regime in Experiment 1, we systematically analyzed how specific parameter manipulations and circuit lesions altered both the ERSPs (Fig. 10) and the corresponding population-specific spike rates across each nucleus (Fig. 11). To isolate intrinsic circuit dynamics from the confounding effects of active postural stabilization, the inverted pendulum was moved passively along predefined periodic trajectories. For this comprehensive spectral analysis, the frequency range of the ERSP was extended up to 100 Hz. For baseline comparison, the ERSPs of the intermittent and forced-choice control models originally shown in Fig. 7 are replotted with this expanded 100-Hz frequency limit in Figs. 10A’ and 10B’, respectively.

**Figure 10:**
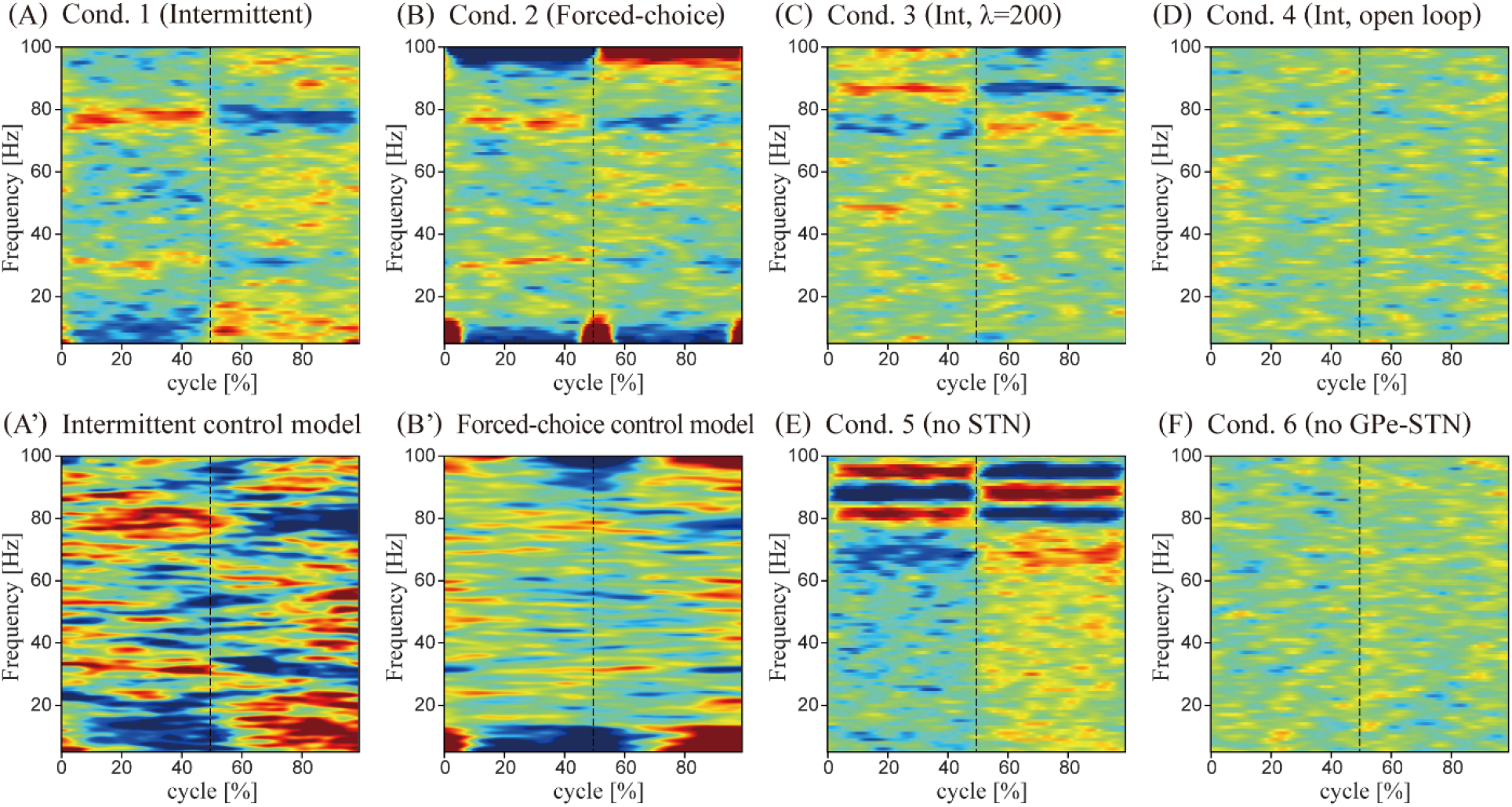
Structural dissections and hyper-sensory drive abolish or alter sway-locked cortical beta modulations. Ensemble averaged ERSP of simulated EEG in Experiment 2 with four conditions. For all conditions, the pendulum moved passively between the regions S_1_ and S_4_ defined in Fig. 2 according to a periodic prescribed trajectory. (A) Condition 1 with the intermittent CBGT controller. (B) Condition 2 with the forced-choice CBGT controller. (C) Condition 3 for a high spike rate of the sensory cortex with the intermittent CBGT controller. (D) Condition 4, where the closed-loop projection from the thalamus to the cortex was severed for the intermittent CBGT controller. (E) Condition 5, where the projection from the cortex to the STN was severed to prevent STN neurons from spiking for the intermittent CBGT controller. (F) Condition 6, where local inhibitory projection from GPe to STN was severed for the intermittent CBGT controller. (A’) and (B’) ERSPs of the intermittent and forced choice control models (Figs. 7A and 7B) are re-displayed, respectively, as (A’) and (B’) with the upper frequency limit extended to 100 Hz.

**Figure 11:**
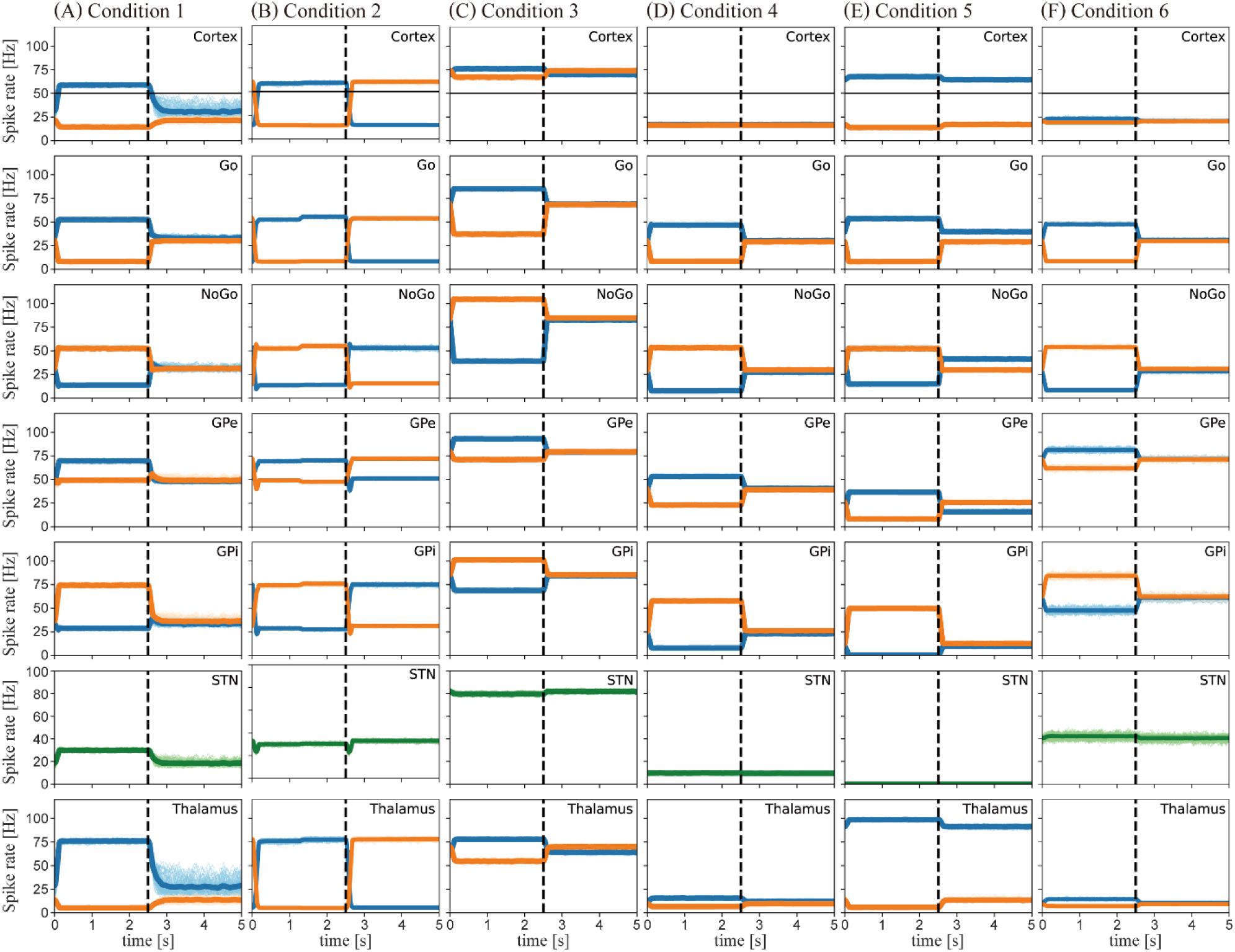
Spike rates of each neuronal population in Experiment 2. From the top to the bottom, spike rates of cortex, Go-striatum, NoGo-striatum, GPe, GPi, STN and Thalamus are shown. Blue and orange lines represent the spike rates of the backward and forward populations, respectively. Thin curves represent the spike rates of 100 single cycles and thick curves represent the spike rates of ensemble average of all cycles. (A) Condition 1. (B) Condition 2. (C) Condition 3. (D) Condition 4, (E) Condition 5, and (F) Condition 6.

First, we examined Condition 1, in which the pendulum was driven along a passive, periodic trajectory designed to mimic a typical sway profile of the intermittent control model, utilizing the original parameter set of the intermittent CBGT controller (Fig. 10A and Fig. 11A). Although the overall modulation amplitudes of ERD and ERS across individual frequency bands were slightly attenuated (Fig. 10A), the resulting ERSP profile was qualitatively identical to that of the active intermittent control model (Fig. 10A’). Furthermore, the movement-cycle dependence of the spike rates in each nucleus (Fig. 11A) remained quantitatively consistent with the active control results (Fig. 8A).

Focusing on the high-frequency range (>50 Hz), which was not detailed in Fig. 7, both Figs. 10A and 10B’ revealed a prominent power modulation around 80 Hz, characterized by an ERS in the first half of the cycle followed by an ERD in the second half. Additionally, weaker power modulations—consisting of an ERD in the first half and an ERS in the second half—were observed in the adjacent bands above (90–100 Hz) and below (50–70 Hz) this central range. Based on the spike-rate profiles (Fig. 11A), the 80-Hz ERS during the first half of the cycle closely corresponds to the aggregate (summed) spike rates, rather than the simple mean, of the backward and forward populations within the cortex and thalamus. Similarly, the subtle ERS in the 50–70 Hz range during the second half of the cycle reflects the aggregate spike rate of these two antagonistic populations during that specific period.

In Condition 2, the pendulum was driven along a passive trajectory mimicking one cycle of the forced-choice control model, utilizing the continuous CBGT parameters (Fig. 10B and Fig. 11B). The beta-band ERSP obtained in Condition 2 closely matched that of the active continuous model (Fig. 10B’), displaying a complete absence of power modulation. Minor discrepancies in the alpha and theta bands between the active and passive conditions were attributable to differences between the unconstrained organic trajectories in Experiment 1 and the prescribed cyclic trajectory in Experiment 2. The underlying mechanism sustaining this tonic, unmodulated beta power can be explained by the stable, tonic spike rates maintained across all nuclei throughout the movement cycle (Fig. 11B), consistent with Fig. 8B.

Intriguingly, in both Figs. 10B and 10B’, phasic modulations emerged in the higher frequency ranges: a distinct 80-Hz rhythm showed an ERS in the first half and an ERD in the second half of the cycle, whereas a 100-Hz rhythm displayed a reversed pattern (an ERD in the first half and an ERS in the second half). Furthermore, smaller-amplitude modulations with an ERS in both the first and second halves emerged around 30 Hz. Crucially, these specific power modulations lacked clear corresponding spike-rate modulations in Fig. 11B, whether evaluating individual, mean, or aggregate population firing rates. This mismatch strongly implies that these high-frequency spectral profiles are emergent phenomena resulting from the intrinsic closed-loop dynamics of the CBGT architecture.

In Condition 3, where the sensory cortex spike rate representing the pendulum state was doubled (*λ*= 200 Hz), the characteristic beta-band power modulation was completely abolished (Fig. 10C and Fig. 11C). The 35-Hz ERS in the first half of the cycle and its subsequent ERD, which were prominent in Condition 1, vanished under this hyper-sensory drive. Instead, the most dominant power modulation shifted to the 80–90 Hz range. This shifted profile consisted of a sub-80-Hz modulation (an ERD in the first half and an ERS in the second half) and adjacent modulations just below and above 80 Hz (characterized by an ERS in the first half and an ERD in the second half, respectively). Although the cortical and thalamic spike rates (Fig. 11C) did not exhibit modulations that directly mirrored these specific frequency boundaries, they appeared to align with the individual population spike rates hovering around 75 Hz, which diverged between the first and second halves of the cycle. However, similar to the findings in Condition 2, these complex spectral signatures represent emergent properties of the closed-loop CBGT circuit dynamics rather than simple linear reflections of firing rates.

In Condition 4, the closed-loop thalamocortical projection was severed in the intermittent CBGT configuration (Fig. 10D and Fig. 11D). Following this disruption, power modulations were completely eliminated across all frequency bands (Fig. 10D). Under this open-loop condition, the cortical spike rates for both the backward and forward populations remained fixed at a tonic baseline of approximately 18 Hz (Fig. 11D). This finding demonstrates that the rhythmic power modulations captured in our simulated EEG are fundamentally driven by the integrity of the broader CBGT closed loop.

In Condition 5, the hyperdirect projection from the cortex to the STN was severed to suppress STN spiking activity within the intermittent CBGT framework (Fig. 10E and Fig. 11E). Under this condition, the ERSP displayed a broad but weak power modulation stretching from 10 to 60 Hz, characterized by a diffuse transition from an ERD in the first half to an ERS in the second half of the cycle. In the high-frequency range, robust power modulations emerged across four distinct, sharply defined bands centered around 70, 80, 90, and 100 Hz. These bands exhibited sequential transitions of ERD-to-ERS, ERS-to-ERD, ERD-to-ERS, and ERS-to-ERD, respectively (Fig. 10E). Because the mean spike rates of the backward and forward populations in both the cortex and thalamus remained constant throughout the movement cycle (Fig. 11E), these spectral shifts cannot be explained by fluctuations in mean firing rates. Instead, these individual high-frequency modulations likely reflect the unmasked firing rates of independent populations encoded within the LFP components. Specifically, the spike rate of the thalamic backward population was approximately 100 Hz in the first half of the cycle and dropped to 90 Hz in the second half, a shift that closely aligns with the ERS observed in those respective bands. Additionally, the prominent 80-Hz ERS during the first half of the cycle appears to be generated by the combined interaction of the cortical and thalamic backward populations, which fired at 60 Hz and 100 Hz, respectively.

Finally, in Condition 6, the local inhibitory feedback projection from the GPe to the STN was severed within the intermittent CBGT controller (Fig. 10F and Fig. 11F). Mirroring the results of the thalamocortical lesion (Condition 4), power modulations were entirely abolished across all frequency bands (Fig. 10F). Cortical spike rates for both populations flattened into a static tonic state at approximately 18 Hz (Fig. 11F). Taken together, these selective dissection results provide definitive evidence that the power modulations characterizing the biomimetic intermittent control model are fundamentally generated by the intrinsic dynamics of the GPe-STN local pacemaker loop.

## Discussion

This study investigated the network-level neural mechanisms underlying sway-phase-locked beta-band modulations during human quiet stance using an embodied spiking neural network (SNN) model of the cortico-basal ganglia-thalamic (CBGT) circuit integrated with a physical inverted pendulum. Our closed-loop simulations demonstrate that the characteristic cortical EEG patterns observed during stance—specifically, beta event-related desynchronization (beta-ERD) during micro-falls and event-related synchronization (beta-ERS / post-movement rebound) during micro-recoveries (Nakamura et al. 2023) —emerge exclusively under an “intermittent control” strategy (*W*_unbiased_). In contrast, a “forced-choice continuous-like” configuration (*W*_biased_), characterized by unremitting active feedback without null-action decision time (DT) intervals, failed to regenerate these dynamic beta modulations. Structural dissections further revealed that these sway-coupled oscillations critically depend on state-dependent sensory feedback, intact thalamocortical loops, and intrinsic STN rhythmicity sustained by reciprocal GPe–STN inhibitory feedback. These findings indicate that CBGT-mediated beta activity serves as a functional neurocomputational biomarker of healthy, intermittent motor selection during postural stabilization.

### Non-circular emergence of beta modulation via embodied body-brain coupling

A primary concern in neuro-computational modeling is whether observed oscillations are artificially hard-coded into model parameters. Our parameter sensitivity analyses strongly refute this “circularity” hypothesis. If beta ERD/ERS were a trivial artifact of *W*_unbiased_, they should have persisted regardless of physical feedback parameters. However, coupling *W*_unbiased_ with large mechanical gains and high noise erased beta modulations. This proves that sway-related beta modulation is not determined by striatal weight topology alone, but is an emergent property generated only when internal CBGT attractor dynamics (balanced Go/NoGo competition permitting prolonging of DT) and physical state dynamics (moderate sway velocity) are mutually matched.

### Physiological plausibility and computational abstractions

While our model uses necessary computational abstractions to execute embodied posture control, its key components are deeply grounded in biological evidence. First, the division of motor cortical and subcortical neurons into forward- and backward-torque populations reflects the somatotopic parallel-channel architecture of the CBGT loop (Alexander and DeLong 1985; Hoover and Strick 1993). Transneuronal tracing confirms discrete representation of opposing muscle groups in primates (Rathelot and Strick 2006), where microstimulation in the putamen evokes discrete single-joint movements (Alexander and DeLong 1985; Nambu et al. 2002). During standing, ankle stabilization relies on antagonist pairs: plantarflexors and dorsiflexors. Modeling competing torque channels thus authentically maps antagonist CBGT motor channels.

Second, while the four-region phase-space partitioning (*S*_1_ − *S*_4_) abstracts multi-sensory integration (somatosensory, vestibular, visual), state-dependent corticostriatal gain modulation is well supported, as the striatum receives dense projections from sensory and motor cortices where synaptic weights dynamically encode state-action mappings. Third, the 50-Hz firing-rate threshold for torque generation aligns with the accumulator/threshold model of movement initiation, supported by primate recordings showing ramping activity crossing neural threshold boundaries prior to movement (Tanaka 2007).

### Physiological mapping of muscle mechanics

Our closed-loop framework aligns directly with the functional division of labor among calf muscles during standing (Loram et al. 2005a; Gawthrop et al. 2011). Background stiffness maintaining ankle stance near equilibrium without excessive energy is provided by tonic Soleus (SOL) activity and intrinsic tendon elasticity (~60–90% of overturning gravitational torque). Conversely, intermittent active feedback torque generated when cortical firing rates cross the decision threshold represents dynamic contractions of Gastrocnemius (MG) and Tibialis Anterior (TA). When sway trajectories reside in safe phase regions (*S*_2_, *S*_4_), active CBGT torque is suspended, leaving the system under intrinsic SOL/tendon stiffness—a state corresponding to prolonged DT and the emergence of beta ERD/ERS.

### Contextualization with Classical Postural Control Frameworks

Continuous sensory-reweighting frameworks (Peterka 2002; Mergner 2010) and intermittent act-and-see models (Loram et al. 2005b; Asai et al. 2009; Gawthrop et al. 2011) have established how sensory integration achieves robust stabilization. Introducing a forced-choice continuous regime within the CBGT network was not designed to refute continuous models, but to serve as a mechanistic contrast. By comparing forced-choice against balanced intermittent regimes, our model demonstrates how spiking CBGT circuits transition between continuous-like torque and discrete intermittent switching, identifying subcortical loop mechanisms that give rise to sway-phase-locked EEG/LFP beta dynamics during human standing.

### Mechanistic synthesis of competing beta modulation hypotheses

Prominent hypotheses regarding beta (13–30 Hz) oscillations include intracortical E/I balance (Jensen et al. 2005; Sherman et al. 2016), thalamocortical resonance (Ching et al. 2010), canonical basal ganglia-thalamo-cortical (BGTC) loop reverberations (Leventhal et al. 2012; Liu et al. 2020), and inter-regional communication maintaining the sensorimotor “status quo” (Bonnefond et al. 2017; Spitzer and Haegens 2017). Furthermore, the GPe–STN reciprocal network acts as a core micro-circuit beta resonator (Koelman and Lowery 2019), whose pathological hypersynchrony characterizes Parkinson’s disease (PD) (Weinberger et al. 2006; Pavlides et al. 2015).

Our model harmonizes these hypotheses within an intermittent motor control architecture. Under intermittent control, STN firing rates modulate phasically based on body physical states. Under continuous control, STN falls into an unmodulated hyper-tonic state (~30 Hz), flattening phase-locked beta rhythmogenesis. Total abolition of beta modulations following hyperdirect pathway (Koketsu et al. 2021) or GPe-STN loop (Fan et al. 2012) lesions confirms that broad BGTC loop integrity is indispensable. Crucially, transient beta dissolution (ERD) and reinstatement (ERS) reflect the computational instantiation of an active intermittent motor-selection loop operating to maintain the sensorimotor status quo.

### Functional balance of sensory corticostriatal projections

The biophysical core driving alternating beta-ERD/ERS resides in the balanced distribution of sensory corticostriatal synaptic weights. Sensory input encodes the instantaneous pendulum state relative to the stable manifold of upright equilibrium. Assigning balanced weights to opposing torque action populations establishes a competitive DT window during which neither motor command gains dominance, manifesting the “control-off” phase of intermittent control. Activity-dependent corticostriatal plasticity (e.g., in dorsolateral striatum) is critical for motor skill refinement (Cataldi et al. 2022), where dopaminergic plasticity alters direct/indirect pathway competition during decision-making (Dunovan et al. 2019). Implementing spike-timing-dependent plasticity (STDP) could enable self-organized acquisition of these balanced profiles.

### Clinical implications and hierarchical postural architecture

Healthy human stance is fundamentally intermittent, whereas postural instability in PD stems from a systemic loss of control intermittency (Perera et al. 2018; Suzuki et al. 2020; Bao and Chen 2024; Bao et al. 2026). This mirrors findings in elderly individuals with chronic dizziness, who display attenuated sway-coupled beta modulations (Ellmers et al. 2025).

While lower-level posture tone and automatic reflexes rely on brainstem (e.g., PPN), cerebellar, and spinal networks (Horak 2006; Takakusaki 2017) —implicitly captured here via data-assimilated physical feedback parameters (Suzuki et al. 2020)—the CBGT loop operates as a higher-order discrete action-selection engine maintaining a “body schema of postural verticality.” Postural impairments in PD (Brown et al. 2001; Jenkinson and Brown 2011; Moisello et al. 2015; Singh 2018; Wu et al. 2019) link directly to this architecture: hyper-stiffness and noise shorten DT, replacing intermittent switching with rigid continuous co-contraction and quenching beta dynamics.

### Methodological limitations, extensibility, and future directions

While leaky integrate-and-fire (LIF) models isolated network topology interactions with physical dynamics, upgrading to conductance-based Hodgkin-Huxley or Izhikevich-type neurons would incorporate intrinsic STN/GPe burst-firing properties. Additionally, driving sensory-corticostriatal weights via autonomous reinforcement learning (Dunovan et al. 2019) could simulate self-organized emergence of postural beta rhythms, since intermittent strategies natively emerge when balancing displacement against metabolic cost (Takazawa et al. 2024). Finally, assimilating our CBGT model with empirical human kinematics and high-density EEG via Bayesian inference (Suzuki et al. 2020; Yokoyama et al. 2025) will provide rigorous data-driven validation.

### A unified neuro-mechanical framework for healthy and Parkinsonian Stance

Traditionally, exaggerated beta oscillations and postural instability in PD have been investigated separately. Our findings provide a unified framework demonstrating that sway-related beta modulations are emergent signatures requiring matched internal CBGT attractor dynamics (DT) and external physical sway dynamics. Applying high mechanical gains and noise characteristic of PD (*P*_large_,*D*_large_, *σ*_large_) quenches beta modulation even under intermittent circuit weights (*W*_unbiased_). Hyper-stiffness forces body states to rapidly traverse safe phase regions (*S*_2_, *S*_4_), depriving CBGT loops of the temporal DT window needed for Go/NoGo competition and collapsing control into a rigid, continuous regime. This reveals how clinical postural phenotypes and electrophysiological biomarkers in PD stem from the same underlying mechanism: the temporal collapse of CBGT decision time driven by altered body–brain loop dynamics.

## Acknowledgments

This study was supported by the Japan Society for the Promotion of Science (JSPS) KAKENHI, No. 22H03662 (T.N.).^1^ The authors sincerely thank Dr. Atsushi Nambu for his insightful discussions and valuable advice on the functional neuroanatomy of the cortico-basal ganglia-thalamic loop and its implementation in our computational model.

## Reference

Alexander GE, DeLong MR (1985) Microstimulation of the primate neostriatum. II. Somatotopic organization of striatal microexcitable zones and their relation to neuronal response properties. J Neurophysiol 53:1417–1430. 10.1152/jn.1985.53.6.1417

Asai Y, Tasaka Y, Nomura K, et al (2009) A Model of Postural Control in Quiet Standing: Robust Compensation of Delay-Induced Instability Using Intermittent Activation of Feedback Control. PLOS ONE 4:e6169. 10.1371/journal.pone.0006169

Bao W, Chen K (2024) Explaining Parkinsonian postural instability using an improved intermittent control model. Chaos Solitons Fractals 182:114844. 10.1016/j.chaos.2024.114844

Bao W, Chen K, Yang Y, et al (2026) Comparative analysis of static and dynamic postural balance control in multiple system atrophy and Parkinson’s disease using an intermittent control model. Gait Posture 127:110167. 10.1016/j.gaitpost.2026.110167

Bevan MD, Magill PJ, Terman D, et al (2002) Move to the rhythm: oscillations in the subthalamic nucleus–external globus pallidus network. Trends Neurosci 25:525–531. 10.1016/S0166-2236(02)02235-X

Bonnefond M, Kastner S, Jensen O (2017) Communication between Brain Areas Based on Nested Oscillations. eNeuro 4:. 10.1523/ENEURO.0153-16.2017

Bottaro A, Yasutake Y, Nomura T, et al (2008) Bounded stability of the quiet standing posture: An intermittent control model. Hum Mov Sci 27:473–495. 10.1016/j.humov.2007.11.005

Brown P, Oliviero A, Mazzone P, et al (2001) Dopamine Dependency of Oscillations between Subthalamic Nucleus and Pallidum in Parkinson’s Disease. J Neurosci 21:1033–1038. 10.1523/JNEUROSCI.21-03-01033.2001

Casadio M, Morasso PG, Sanguineti V (2005) Direct measurement of ankle stiffness during quiet standing: implications for control modelling and clinical application. Gait Posture 21:410–424. 10.1016/j.gaitpost.2004.05.005

Cataldi S, Stanley AT, Miniaci MC, Sulzer D (2022) Interpreting the role of the striatum during multiple phases of motor learning. FEBS J 289:2263–2281. 10.1111/febs.15908

Ching S, Cimenser A, Purdon PL, et al (2010) Thalamocortical model for a propofol-induced α-rhythm associated with loss of consciousness. Proc Natl Acad Sci 107:22665–22670. 10.1073/pnas.1017069108

Dunovan K, Vich C, Clapp M, et al (2019) Reward-driven changes in striatal pathway competition shape evidence evaluation in decision-making. PLOS Comput Biol 15:e1006998. 10.1371/journal.pcbi.1006998

Ellmers TJ, Ibitoye R, Castro P, et al (2025) Chronic dizziness in older adults: Disrupted sensorimotor EEG beta oscillations during postural instability. Clin Neurophysiol 174:31–36. 10.1016/j.clinph.2025.03.032

Fan KY, Baufreton J, Surmeier DJ, et al (2012) Proliferation of External Globus Pallidus-Subthalamic Nucleus Synapses following Degeneration of Midbrain Dopamine Neurons. J Neurosci 32:13718–13728. 10.1523/JNEUROSCI.5750-11.2012

Gawthrop P, Loram I, Lakie M, Gollee H (2011) Intermittent control: a computational theory of human control. Biol Cybern 104:31–51. 10.1007/s00422-010-0416-4

Holgado AJN, Terry JR, Bogacz R (2010) Conditions for the Generation of Beta Oscillations in the Subthalamic Nucleus–Globus Pallidus Network. J Neurosci 30:12340–12352. 10.1523/JNEUROSCI.0817-10.2010

Hoover JE, Strick PL (1993) Multiple Output Channels in the Basal Ganglia. Science 259:819–821. 10.1126/science.7679223

Horak FB (2006) Postural orientation and equilibrium: what do we need to know about neural control of balance to prevent falls? Age Ageing 35:ii7–ii11. 10.1093/ageing/afl077

Izhikevich EM (2004) Which model to use for cortical spiking neurons? IEEE Trans Neural Netw 15:1063–1070. 10.1109/TNN.2004.832719

Jenkinson N, Brown P (2011) New insights into the relationship between dopamine, beta oscillations and motor function. Trends Neurosci 34:611–618. 10.1016/j.tins.2011.09.003

Jensen O, Goel P, Kopell N, et al (2005) On the human sensorimotor-cortex beta rhythm: Sources and modeling. NeuroImage 26:347–355. 10.1016/j.neuroimage.2005.02.008

Johnson DH (1996) Point process models of single-neuron discharges. J Comput Neurosci 3:275–299. 10.1007/BF00161089

Koelman LA, Lowery MM (2019) Beta-Band Resonance and Intrinsic Oscillations in a Biophysically Detailed Model of the Subthalamic Nucleus-Globus Pallidus Network. Front Comput Neurosci 13:. 10.3389/fncom.2019.00077

Koketsu D, Chiken S, Hisatsune T, et al (2021) Elimination of the Cortico-Subthalamic Hyperdirect Pathway Induces Motor Hyperactivity in Mice. J Neurosci 41:5502–5510. 10.1523/JNEUROSCI.1330-20.2021

Leventhal DK, Gage GJ, Schmidt R, et al (2012) Basal Ganglia Beta Oscillations Accompany Cue Utilization. Neuron 73:523–536. 10.1016/j.neuron.2011.11.032

Lindi SA, Mallet NP, Leblois A (2024) Synaptic Changes in Pallidostriatal Circuits Observed in the Parkinsonian Model Triggers Abnormal Beta Synchrony with Accurate Spatio-temporal Properties across the Basal Ganglia. J Neurosci 44:. 10.1523/JNEUROSCI.0419-23.2023

Liu C, Zhou C, Wang J, et al (2020) The role of coupling connections in a model of the cortico-basal ganglia-thalamocortical neural loop for the generation of beta oscillations. Neural Netw 123:381–392. 10.1016/j.neunet.2019.12.021

Loram ID, Lakie M (2002) Direct measurement of human ankle stiffness during quiet standing: the intrinsic mechanical stiffness is insufficient for stability. J Physiol 545:1041–1053. 10.1113/jphysiol.2002.025049

Loram ID, Maganaris CN, Lakie M (2005a) Active, non-spring-like muscle movements in human postural sway: how might paradoxical changes in muscle length be produced? J Physiol 564:281–293. 10.1113/jphysiol.2004.073437

Loram ID, Maganaris CN, Lakie M (2005b) Human postural sway results from frequent, ballistic bias impulses by soleus and gastrocnemius. J Physiol 564:295–311. 10.1113/jphysiol.2004.076307

McCarthy MM, Moore-Kochlacs C, Gu X, et al (2011) Striatal origin of the pathologic beta oscillations in Parkinson’s disease. Proc Natl Acad Sci 108:11620–11625. 10.1073/pnas.1107748108

Mergner T (2010) A neurological view on reactive human stance control. Annu Rev Control 34:177–198. 10.1016/j.arcontrol.2010.08.001

Mohanty B, Guo Z, Johnson LA, et al (2025) Parkinsonism disrupts the balance between excitatory and inhibitory activity within the primary motor cortex during movement. Proc Natl Acad Sci 122:e2510287122. 10.1073/pnas.2510287122

Moisello C, Blanco D, Lin J, et al (2015) Practice changes beta power at rest and its modulation during movement in healthy subjects but not in patients with Parkinson’s disease. Brain Behav 5:e00374. 10.1002/brb3.374

Nakamura A, Miura R, Suzuki Y, et al (2023) Discrete cortical control during quiet stance revealed by desynchronization and rebound of beta oscillations. Neurosci Lett 814:137443. 10.1016/j.neulet.2023.137443

Nambu A, Kaneda K, Tokuno H, Takada M (2002) Organization of Corticostriatal Motor Inputs in Monkey Putamen. J Neurophysiol 88:1830–1842. 10.1152/jn.2002.88.4.1830

Neuper C, Pfurtscheller G (2001) Evidence for distinct beta resonance frequencies in human EEG related to specific sensorimotor cortical areas. Clin Neurophysiol 112:2084–2097. 10.1016/S1388-2457(01)00661-7

Nomura T, Suzuki Y, Morasso PG (2022) Intermittent Control Strategy for Stabilizing Human Quiet Stance, A Model of the. In: Encyclopedia of Computational Neuroscience. Springer, New York, NY, pp 1694–1704

Ouyang G, Wang S, Liu M, et al (2022) Multilevel and multifaceted brain response features in spiking, ERP and ERD: experimental observation and simultaneous generation in a neuronal network model with excitation–inhibition balance. Cogn Neurodyn. 10.1007/s11571-022-09889-w

Pavlides A, Hogan SJ, Bogacz R (2015) Computational Models Describing Possible Mechanisms for Generation of Excessive Beta Oscillations in Parkinson’s Disease. PLOS Comput Biol 11:e1004609. 10.1371/journal.pcbi.1004609

Perera T, Tan JL, Cole MH, et al (2018) Balance control systems in Parkinson’s disease and the impact of pedunculopontine area stimulation. Brain 141:3009–3022. 10.1093/brain/awy216

Peterka RJ (2002) Sensorimotor Integration in Human Postural Control. J Neurophysiol 88:1097–1118. 10.1152/jn.2002.88.3.1097

Pfurtscheller G, Lopes da Silva FH (1999) Event-related EEG/MEG synchronization and desynchronization: basic principles. Clin Neurophysiol 110:1842–1857. 10.1016/S1388-2457(99)00141-8

Ratcliff R, Smith PL, Brown SD, McKoon G (2016) Diffusion Decision Model: Current Issues and History. Trends Cogn Sci 20:260–281. 10.1016/j.tics.2016.01.007

Rathelot J-A, Strick PL (2006) Muscle representation in the macaque motor cortex: An anatomical perspective. Proc Natl Acad Sci 103:8257–8262. 10.1073/pnas.0602933103

Roopun AK, Kramer MA, Carracedo LM, et al (2008) Period concatenation underlies interactions between gamma and beta rhythms in neocortex. Front Cell Neurosci 2:. 10.3389/neuro.03.001.2008

Roopun AK, LeBeau FE, Rammell J, et al (2010) Cholinergic neuromodulation controls directed temporal communication in neocortex in vitro. Front Neural Circuits 4:. 10.3389/fncir.2010.00008

Sherman MA, Lee S, Law R, et al (2016) Neural mechanisms of transient neocortical beta rhythms: Converging evidence from humans, computational modeling, monkeys, and mice. Proc Natl Acad Sci 113:E4885–E4894. 10.1073/pnas.1604135113

Singh A (2018) Oscillatory activity in the cortico-basal ganglia-thalamic neural circuits in Parkinson’s disease. Eur J Neurosci 48:2869–2878. 10.1111/ejn.13853

Spitzer B, Haegens S (2017) Beyond the Status Quo: A Role for Beta Oscillations in Endogenous Content (Re)Activation. eNeuro 4:. 10.1523/ENEURO.0170-17.2017

Stančák A, Feige B, Lücking CH, Kristeva-Feige R (2000) Oscillatory cortical activity and movement-related potentials in proximal and distal movements. Clin Neurophysiol 111:636–650. 10.1016/S1388-2457(99)00310-7

Steriade M (2006) Grouping of brain rhythms in corticothalamic systems. Neuroscience 137:1087–1106. 10.1016/j.neuroscience.2005.10.029

Suzuki Y, Nakamura A, Milosevic M, et al (2020) Postural instability via a loss of intermittent control in elderly and patients with Parkinson’s disease: A model-based and data-driven approach. Chaos Interdiscip J Nonlinear Sci 30:113140. 10.1063/5.0022319

Tachibana Y, Iwamuro H, Kita H, et al (2011) Subthalamo-pallidal interactions underlying parkinsonian neuronal oscillations in the primate basal ganglia. Eur J Neurosci 34:1470–1484. 10.1111/j.1460-9568.2011.07865.x

Takakusaki K (2017) Functional Neuroanatomy for Posture and Gait Control. J Mov Disord 10:1–17. 10.14802/jmd.16062

Takazawa T, Suzuki Y, Nakamura A, et al (2024) How the brain can be trained to achieve an intermittent control strategy for stabilizing quiet stance by means of reinforcement learning. Biol Cybern 118:229–248. 10.1007/s00422-024-00993-0

Tanaka M (2007) Cognitive Signals in the Primate Motor Thalamus Predict Saccade Timing. J Neurosci 27:12109–12118. 10.1523/JNEUROSCI.1873-07.2007

Terman D, Rubin JE, Yew AC, Wilson CJ (2002) Activity Patterns in a Model for the Subthalamopallidal Network of the Basal Ganglia. J Neurosci 22:2963–2976. 10.1523/JNEUROSCI.22-07-02963.2002

Ursino M, Véronneau-Veilleux F, Nekka F (2020) A non-linear deterministic model of action selection in the basal ganglia to simulate motor fluctuations in Parkinson’s disease. Chaos Interdiscip J Nonlinear Sci 30:083139. 10.1063/5.0013666

Vinding MC, Tsitsi P, Piitulainen H, et al (2019) Attenuated beta rebound to proprioceptive afferent feedback in Parkinson’s disease. Sci Rep 9:2604. 10.1038/s41598-019-39204-3

Weinberger M, Mahant N, Hutchison WD, et al (2006) Beta Oscillatory Activity in the Subthalamic Nucleus and Its Relation to Dopaminergic Response in Parkinson’s Disease. J Neurophysiol 96:3248–3256. 10.1152/jn.00697.2006

Winter DA, Patla AE, Prince F, et al (1998) Stiffness Control of Balance in Quiet Standing. J Neurophysiol 80:1211–1221. 10.1152/jn.1998.80.3.1211

Wu H-M, Hsiao F-J, Chen R-S, et al (2019) Attenuated NoGo-related beta desynchronisation and synchronisation in Parkinson’s disease revealed by magnetoencephalographic recording. Sci Rep 9:7235. 10.1038/s41598-019-43762-x

Yokoyama H, Noda Y, Wada M, et al (2025) Validation of an electroencephalography data assimilation-based computational approach for estimating cortical excitation-inhibition balance. Commun Eng 4:195. 10.1038/s44172-025-00525-z

